# Quantifying the effects of intraspecific trait variation and interspecific trait correlations on interacting populations - a nonlinear averaging approach

**DOI:** 10.1101/2021.03.11.435001

**Authors:** Koen J. van Benthem, Rishabh Bagawade, Chantal Blüml, Peter Nabutanyi, Frans M. Thon, Meike J. Wittmann

## Abstract

Interactions between two species, e.g. between a predator species and a prey species, can often be described as the sum of many individual-by-individual interactions whose outcomes depend on the traits of the interacting individuals. These traits often vary substantially among individuals in each species, and individuals do not always interact randomly, e.g. due to plastic responses to a shared environmental factor in a heterogeneous landscape. Here we investigate the impact of intraspecific trait variation (ITV) and such interspecific trait correlations on species interactions via nonlinear averaging. Building on past models that integrate over an interaction kernel to obtain the impacts of ITV, we develop a modeling framework that allows to model arbitrary species interactions, with interspecific trait correlations as novel feature. Based on two key ingredients, a joint trait distribution and a two-dimensional interaction function, the average interaction parameters (e.g. average predation rate) can be quantified numerically, approximated using an insightful Taylor approximation, and compared to cases without ITV. We highlight two applications of our framework. First, we study the quantitative and qualitative effects of ITV and trait correlations in a simple predator-prey model and show that even in the absence of evolution, variation and trait correlations among interacting individuals can make or break the coexistence between species. Second, we use simulated field data for a predator-prey system to show how the impact of ITV on an ecological interaction can be estimated from empirical data.

**Highlights:**

- We model how intraspecific trait variation in two species affects their interaction.
- Our framework allows correlations between the traits of interacting individuals.
- Trait variation and correlations can strongly affect ecological interactions.
- Trait variation and correlations can make or break coexistence.
- The effect of intraspecific trait variation can be estimated from data.

## 1 Introduction

Intraspecific trait variation (ITV) is increasingly being recognized as a factor that can considerably affect population dynamics, community and ecosystem patterns, and global change responses (Bolnick et al., 2011; Des Roches et al., 2018; Moran et al., 2016; Raffard et al., 2019; Violle et al., 2012). For example, ITV can affect population growth of a single species (Bjørnstad and Hansen, 1994), influence whether or not competing species can coexist (e.g. Hart et al., 2016; Hausch et al., 2018; Holdridge and Vasseur, 2022; Maynard et al., 2019; Stump et al., 2022; Uriarte and Menge, 2018), affect predator-prey dynamics (Coblentz et al., 2021; Pettorelli et al., 2011, 2015), and even affect the qualitative outcome of a biological interaction (Moran et al., 2022), i.e. whether it is antagonistic or cooperative.

Here we focus on cases where two species engage in direct individual-by-individual interactions whose outcome depends on the exact trait combinations of the interacting individuals. Examples of such interactions include the performance of a plant depending on its own genotype and traits and those of neighbors from a competing species (Fridley et al., 2007; Genung et al., 2012), the ability of a predator to consume a prey depending on the ratio of predator mass to prey mass (Nakazawa, 2017; Portalier et al., 2019), and the ability of an insect to pollinate a flower depending on the size of the insect’s proboscis relative to the length and width of the flower’s corolla (Ibanez, 2012). Furthermore, whether or not individuals interact at all can be nonrandom and trait-dependent. For example, trait-specific preferences for certain microhabitats are expected to induce positive trait correlations between the boldness of interacting individuals in bank voles and striped field mice (Schirmer et al., 2020) and between activity levels in interacting individuals of different stickleback species (Webster et al., 2008). Trait correlations are also expected to emerge when predators only search for prey that is within a feasible range given their own body size (Portalier et al., 2019), or in plants in response to a common elevation gradient (Halbritter et al., 2018). Focusing on species mean traits to describe such interactions could lead to misleading results, for example, a naive conclusion that individuals of two species cannot interact based on average trait values, when in fact interactions between many pairs of individuals are possible (González-Varo and Traveset, 2016). This is why it has been argued that the focus of community ecology should shift from species-species interactions to individual-individual interactions, in the same way there has been a shift from the species to the individual as the unit of selection in evolutionary biology (Nakazawa, 2017).

One important mechanism by which ITV influences population dynamics is nonlinear averaging: when a demographic parameter, e.g. reproductive rate, is a nonlinear function of a certain trait, the average value of the demographic parameter across individuals is in general not equal to the demographic parameter of an individual with average trait value (Bjørnstad and Hansen, 1994; Ruel and Ayres, 1999). In such cases, ITV can lead to directional changes in population-level properties. The direction and magnitude of the effect depend on the curvature of the function relating the trait and the demographic parameter, according to Jensen’s inequality (Jensen, 1905, see section 2.2.1 for more detailed explanations). Nonlinear averaging has been taken into account in some explorations of the conditions under which ITV promotes or hinders the coexistence of competitors (Gravel et al., 2011; Hart et al., 2016; Stump et al., 2022; Uriarte and Menge, 2018) or affects predator-prey dynamics (Gibert and Brassil, 2014; Okuyama, 2008; Schreiber et al., 2011).

Some of these studies assume that there is ITV in only one species. For example, Gibert and Brassil (2014) explored the consequences of ITV in predator attack rate and handling time for predation efficiency in a Rosenzweig-MacArthur predator-prey model. They used bell-shaped functions to describe predation efficiency, with the highest predation efficiency at an optimum intermediate predator trait. In other studies, there is ITV in two or more species, but each species is only indirectly affected by the ITV in the other species, e.g. via population density (e.g. results on variation in competitive ability in Hart et al., 2016) or shared resources (Holdridge and Vasseur, 2022), so that it is sufficient to deal with ITV in one species at a time. In addition, there is limited research investigating the effects of ITV in two or more species where the outcome of interaction depends on the exact trait combination of interacting individuals. Such ITV can be taken into account by integrating over interaction kernels, i.e. functions that state the probability or outcome of an interaction as a function of both interaction partners’ trait values (e.g. Barabás and D’Andrea, 2016; Hart et al., 2016; McPeek et al., 2022; Senthilnathan and Gavrilets, 2021).

Here we develop a general nonlinear averaging framework to understand the population-level effects of joint trait variation in two species engaged in direct individual-by-individual interactions. Our framework addresses two issues with previous work: First, the possibility that individuals may encounter each other nonrandomly is not included in most previous work. To better understand the effects of ITV, we thus need to account for trait correlations across multiple species. Second, while most previous research on the effects of ITV with individual-by-individual interactions has been on interspecific competition (but see Senthilnathan and Gavrilets, 2021), our framework generalizes to all types of species interactions described by arbitrary interaction functions. In addition to a numerical approach, we provide a flexible analytic approximation that provides an intuitive understanding of the effects of ITV and correlations in two interacting species. In two application examples, we then highlight two ways in which our framework can be useful: first, we show in the example of a predator-prey system how ITV and interspecific trait correlations can lead to quantitative and even qualitative changes in population dynamics and coexistence outcomes. Second, we use simulated data to show how our framework could be applied to estimate the effects of ITV on average interaction parameters from empirical data.

## 2 General modeling framework

### 2.1 Joint trait distribution and interaction function

We focus on a system of two interacting species *A* and *B* characterized by mean traits *µ*_*a*_ and *µ*_*b*_, trait variances 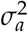 and 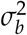 and trait correlation *ρ*_*a,b*_ between interacting individuals. Our ap-proach has two key ingredients (figure 1, left): 1) a bivariate joint trait distribution *f*_*AB*_(*a, b*) that models for a pair of interacting individuals how likely the species-*A* individual is to have trait value *a* and the species-*B* individual trait value *b*. We assume this distribution to be a bivariate normal distribution throughout the manuscript. However, our general framework can be applied to any bivariate distribution that stays constant over the ecological time scale of interest. 2) any interaction function *γ*(*a, b*) that gives the rate and outcome of the pairwise interaction between individuals with trait values *a* and *b*. To allow for analytic approximations, the interaction func-tion should be at least twice differentiable with respect to both trait values.

**Figure 1.**
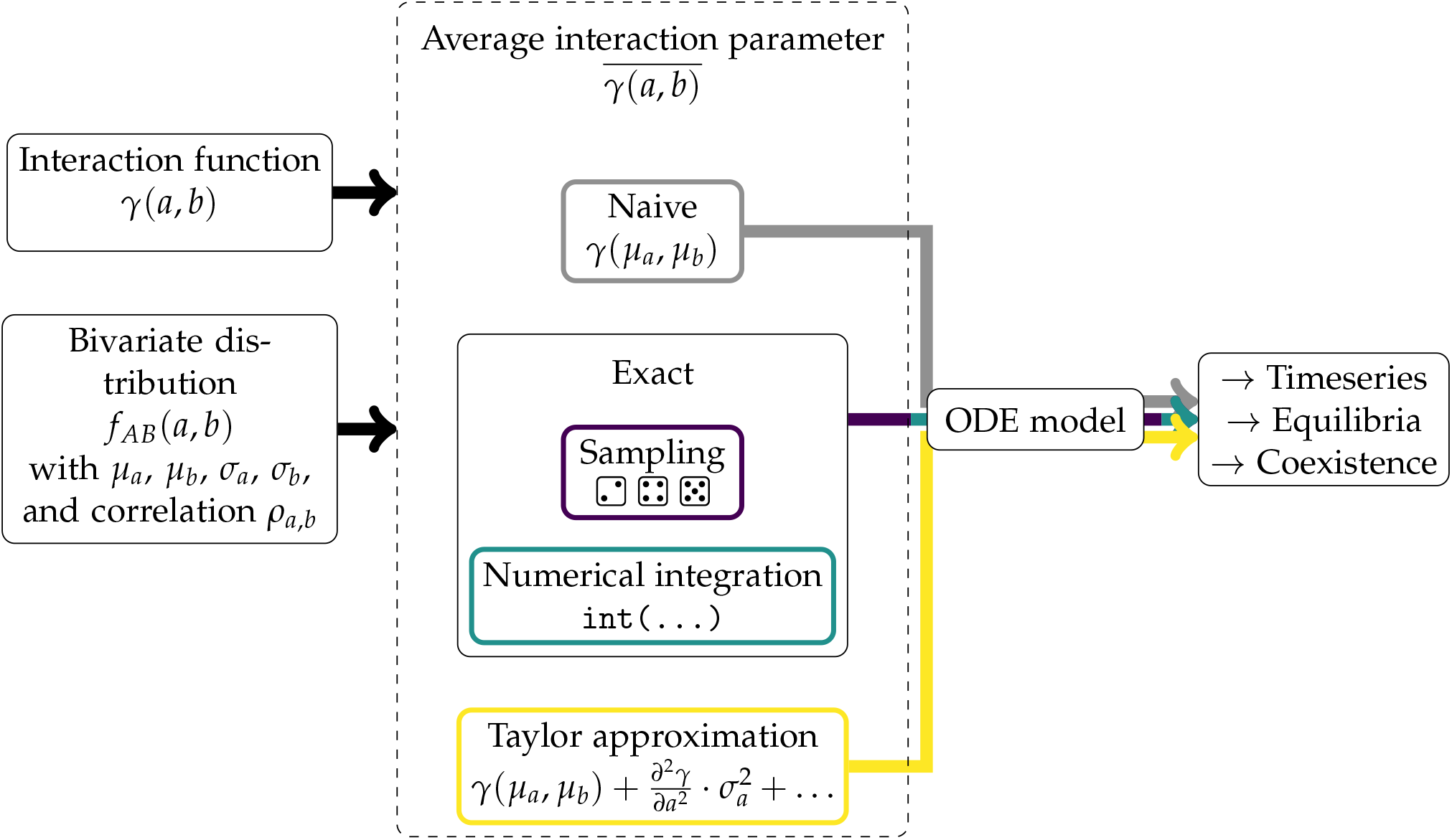
Overview of the general modeling approach. Each analysis starts with an interaction function, *γ*, that describes the rate and outcome of the interaction between individuals with trait value *a* and *b* and a joint trait distribution for pairs of interacting individuals from species *A* and species *B*. In the examples that we show, we use a bivariate normal distribution that is characterised by the trait means of both species (*µ*_*a*_ and *µ*_*b*_), the standard deviations of these trait values (*σ*_*a*_ an *σ*_*b*_) and the interspecific trait correlation (*ρ*_*a,b*_). From these two functions, the average interaction parameter is calculated using three different methods (a naive approach, and two forms of nonlinear averaging: sampling for obtaining the exact effect size and a Taylor approximation). Another way of computing the exact value via numerical integration is shown in S1. Finally, each estimated value for the average interaction parameter can be substituted into an ordinary differential equation (ODE) model to predict the resulting differences in population dynamics. R code and a shiny app to run this workflow for a given interaction function and joint trait distribution are provided in the online supporting information.

The interaction function can be estimated from empirical data (Bartomeus et al., 2016), either by fitting a curve with a predefined shape, or by fitting a smooth function without specifying a shape beforehand (Wetzel et al., 2016; Wood, 2006). We use the latter method in section 5 below on simulated data to illustrate how the nonlinear averaging effect of joint ITV can be estimated from empirical data.

#### 2.1.1 Example for the mechanistic underpinning of a joint trait distribution

Let us now consider one example of how a joint trait distribution can emerge mechanistically. Assume that individuals of both species inhabit patches (shared with a number of individuals both of the same and of the other species) and that their traits respond plastically to the micro-environmental conditions in their patch. In addition, there is some additional individual-specific random effect on trait values. The trait value of an individual *k* of species *i* in patch *j* is then given by

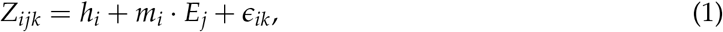

where *h*_*i*_ and *m*_*i*_ are the intercept and slope of the reaction norm of species *i, E*_*j*_ is the micro-environment in patch *j* which we here assume to have mean 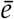 and variance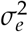, and ϵ_*ik*_ is a random individual trait effect that is drawn from an independent normal distribution with mean 0 and variance 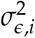. We assume that individuals can only interact with other individuals in the same patch and that individuals are distributed uniformly over patches so that the total rate at which individuals co-occur with others is the same for all individuals and independent of traits. The joint trait distribution of interacting individuals (i.e. those in the same patch) is then characterized by

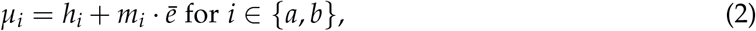

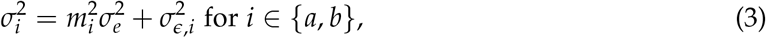

and

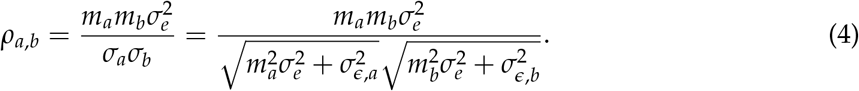

Note that if the mean of the environmental values changes, e.g. due to climate change, means of the marginal trait distributions will also change, but variances and trait correlations will remain constant in this model.

### 2.2 Quantifying the average interaction parameter

To determine the ecological consequences of trait variation in the interaction partners, we first determine the average outcome 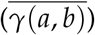 of an interaction across all possible pairs of individuals, weighted by the probability of this pairing given the joint trait distribution *f*_*AB*_(*a, b*) (figure 1):

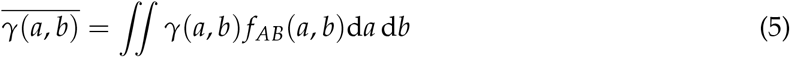

This assumes that the population-level outcome of the interaction can be thought of as the sum of many individual-by-individual interactions.

We solve the double integral in Eq. (5) by averaging the interaction function over 500,000 trait-value pairs randomly sampled from the joint trait distribution in R (R Core Team, 2022). This sample size led to acceptably small sampling noise, as indicated by visually nearly indistinguishable results from other numerical integration methods (described in S1). In the following, we refer to the sampling value as the exact value of the average interaction parameter, keeping in mind its small sampling error.

We compare the average value of *γ*(*a, b*), 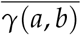, to a naive estimate of the interaction strength, *γ*(*µ*_*a*_, *µ*_*b*_), which is based only on the species mean trait values and is equivalent to a case without ITV.

#### 2.2.1 Analytic approximation

Next, we derive an insightful analytic approximation for the average interaction parameter with ITV in both interacting species. We first briefly describe the state of the art for ITV in a single species and then extend the approximation to two dimensions, one for each species.

Based on Jensen’s inequality (Jensen, 1905), the direction of the effect of ITV on the average value of a nonlinear function depends on the curvature of the nonlinear function (figure 2A). One can measure this curvature through the second derivative. If the function has a positive second derivative (i.e. is convex) around the species mean trait (around the black point in figure 2A), a small amount of intraspecific trait variation will lead to an increase in the average value of the demographic parameter; and if the function has a negative second derivative around the species mean trait (i.e. is concave, around the grey point in figure 2A), a small amount of ITV will reduce the average value of the demographic parameter. Quantifying the second derivative of an ecological response variable with respect to some trait value can thus give insight into the consequences of ITV (Inouye, 2005). For example, Wetzel et al. (2016) found a negative second derivative in the function linking plant nutrient levels to herbivore performance, leading to a negative effect of variation in nutrient levels on herbivore performance. By contrast, in predator species with a type 2 functional response, consumption rate as a function of handling time has a positive second derivative such that individual variation in handling time increases average predator consumption rate (Bolnick et al., 2011).

**Figure 2.**
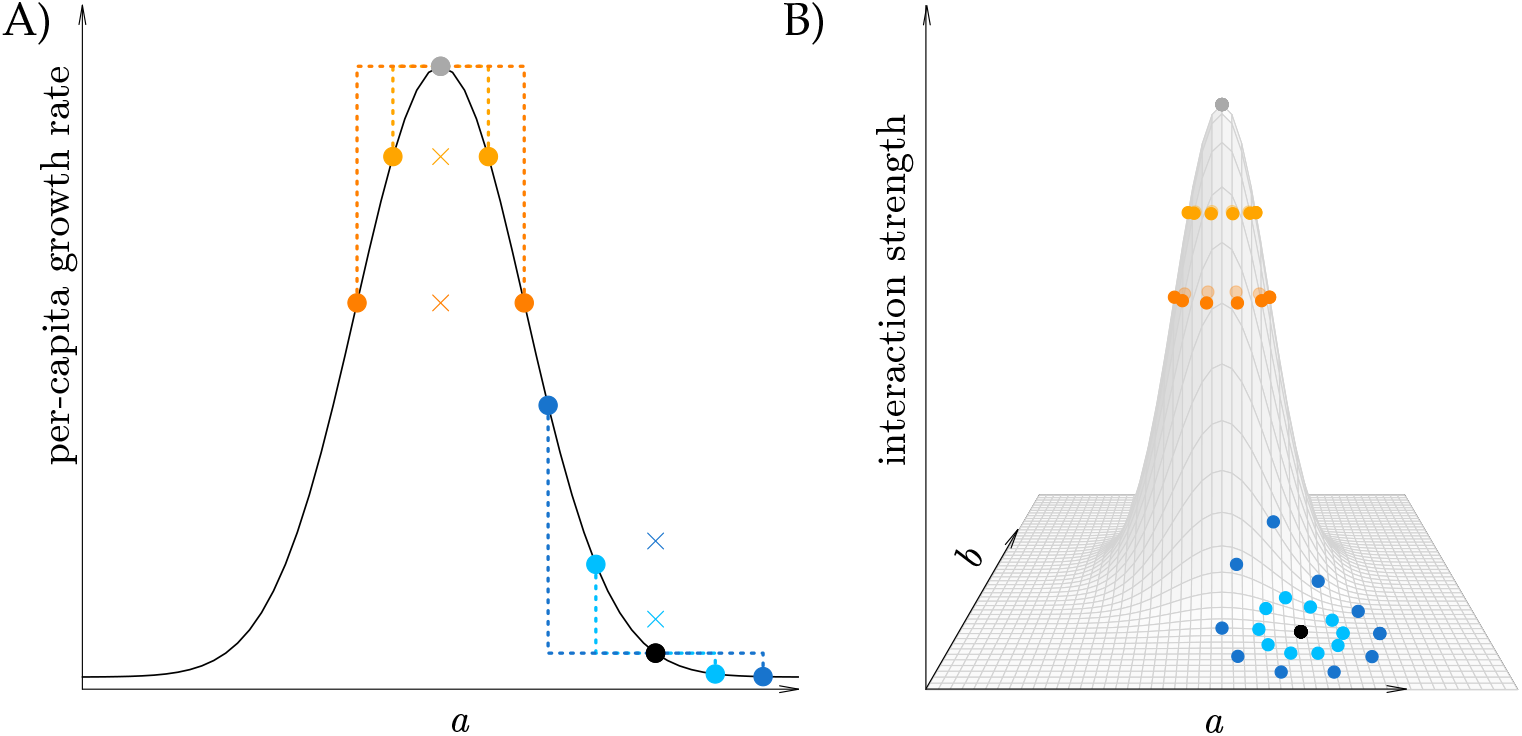
Illustration of nonlinear averaging in one and two dimensions. A) A demographic property such as per-capita growth rate depends on trait values *a*. In a population with a mean trait value near the maximum of a Gaussian function (grey point), the further two *a* values are spread around this mean, the more their average per-capita growth rate (crosses) decreases (compare orange and yellow crosses to the grey dot). This can be explained by the negative curvature around the maximum. By contrast, in a population with mean trait value close to the base of the curve (black point), the function has a positive curvature and therefore the opposite occurs: the average per-capita growth rate (blue crosses) for two blue points that are equally far from the black, middle point, is larger than the per-capita growth rate at the average trait value (black point). B) In two dimensions, when trait values *a* and *b* vary in two interacting species, a similar mechanism may affect the average interaction strength: when trait means lie around the maximum of a bell-shaped interaction function (grey dot), the curvatures are negative in all directions and as points are further spread in the *a, b* − plane around the maximum, their average height decreases (orange dots). When the trait means are close towards the base of the function (black dot), the curvatures are positive, and the opposite effect occurs: here an increase in variation in *a* and *b* (from the black to the dark blue points) leads to an increase in the average interaction strength, which may represent for example competition strength or predation rate.

The effect of ITV on the average value of a nonlinear function works very similarly in two dimensions (figure 2B) and this can be captured in an analytic approximation based on a second-order Taylor series around the point (*µ*_*a*_, *µ*_*b*_) (see S2.1 for derivation):

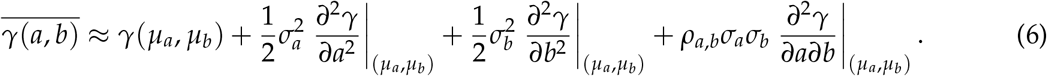

That is, ITV in species A has a positive or negative effect depending on whether the second derivative of the interaction function with respect to the trait of the species-A individual is positive or negative, and analogously for species B. Moreover, the direction and magnitude of the effect of interspecific trait correlations depends on the mixed second derivative. To evaluate the derivatives in Eq. (6) for any interaction function, we use the Deriv package in R (Clausen and Sokol, 2020).

Note that similar approximations have been used previously to understand the effects of variation in two potentially correlated environmental factors (Koussoroplis et al., 2017) and in two potentially correlated traits (Koussoroplis et al., 2019) on population parameters. When traits are symmetrically distributed, the third order errors are zero (S2.2) and the magnitude of the error of this approximation is therefore of fourth order in deviations from the respective mean traits. Thus, the approximation will fit well as long as most trait values are close to the mean, but might break down for large trait variances. Because third order errors do not disappear for asymmetric distributions, we expect that in such cases the accuracy will go down more quickly with increasing trait variances. The evaluation of this Taylor approximation is generally less computationally intensive compared to the numerical integration. However, the main benefit is that the approximation makes it easier to grasp intuitively how the effect of ITV and trait correlation on the average interaction function is determined by the shape of the interaction function.

### 2.3 Illustration using two interaction functions

We now illustrate the three described estimation methods (see figure 1) on a general second-order two-dimensional polynomial and a logistic interaction function (Table 1). For each function, we determine the magnitude and direction of the effect of ITV and trait correlation on the average interaction value, as well as the accuracy of the approximation. These examples serve as illustrations, but note that the approach is flexible and not limited to a bivariate normal joint trait distribution, nor to the used interaction functions.

**Table 1.** Overview of the example interaction functions and their second-order partial derivatives used in Eq. 6 to approximate 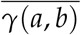.

| Interaction function | $\gamma(a, b)$ | $\frac{\partial^2 \gamma}{\partial a^2}$ | $\frac{\partial^2 \gamma}{\partial b^2}$ | $\frac{\partial^2 \gamma}{\partial a \partial b}$ |
| --- | --- | --- | --- | --- |
| Polynomial | $p_1 a^2 + p_2 b^2 + p_3 ab + p_4 a + p_5 b + p_6$ | $2p_1$ | $2p_2$ | $p_3$ |
| Logistic | $p_{\max} \frac{1}{1 + e^{-k(b-a-h)}}$ | $k^2(\gamma(a, b) + \frac{2\gamma(a, b)^3}{p_{\max}^2} - 3\frac{\gamma(a, b)^2}{p_{\max}})$ | $\frac{\partial^2 \gamma}{\partial a^2}$ | $-\frac{\partial^2 \gamma}{\partial a^2}$ |

The first interaction function we explore is a *two-dimensional polynomial of order two*:

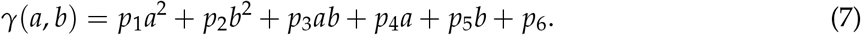

We chose this interaction function because of its flexible shape. Depending on the parameter values (*p*_*i*_), this interaction function is shaped as a plane, paraboloid, or a saddle. Since the sign of the interaction can change with the trait values, it can for example be used to describe predator-prey reversal. However, the interaction function is not bounded, and when using it, one should verify that it takes reasonable values over the range of trait values considered. Alternatively, one might want to introduce bounds (see S3.2 for an example).

Using the partial derivatives from Table 1, we obtain an approximation for the average interaction parameter, which is exact since the third and higher order derivatives are zero for this interaction function (see also S2.2):

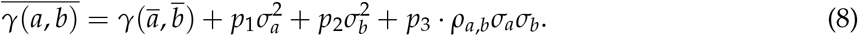

If, for example, *p*_1_, *p*_2_, and *p*_3_ are zero, the interaction function reduces to a linear function in *a* and *b*, and intraspecific variation has no effect on 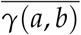.

If instead *p*_1_, *p*_2_, and *p*_3_ are all negative, with *p*_3_ having the largest absolute value (figure 3A), the effect of variation in *a* and *b* on the average interaction function 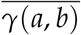 is negative when *ρ*_*a,b*_ is zero or close to zero (figure 3 green ellipses in A and green bar in B). This is caused by the negative curvature of the function in the *a* and *b* directions. The negative effect is even stronger with a positive interspecific trait correlation (light orange and pink scenarios in figure 3 A,B). Along the line *a* = −*b*, in contrast, the curvature is positive, leading to a positive effect of trait variation on the average interaction parameter when the individual trait value combinations fall along this line, i.e. when trait correlation is strongly negative (figure 3 A-B, dark orange scenario, similarly in the blue scenario). Whether the trait correlation can change the sign of the effect of ITV depends on the value of *p*_3_ relative to *p*_1_ and *p*_2_ (section S3.1). Biologically, an interaction function of this general shape could for example represent a mutualism that only works when interacting individuals have diversified (complementary) traits, but turns into competition when traits are similar. Interestingly, although the magnitude of the interaction changes with the mean trait values (compare the left two bars in figure 3B to the right three), the difference between the naive value and the other estimates only depends on *σ*_*a*_, *σ*_*b*_ and *ρ*_*a,b*_ (figure 3B, the effects of ITV are the same e.g. in the blue and dark orange scenario).

**Figure 3.**
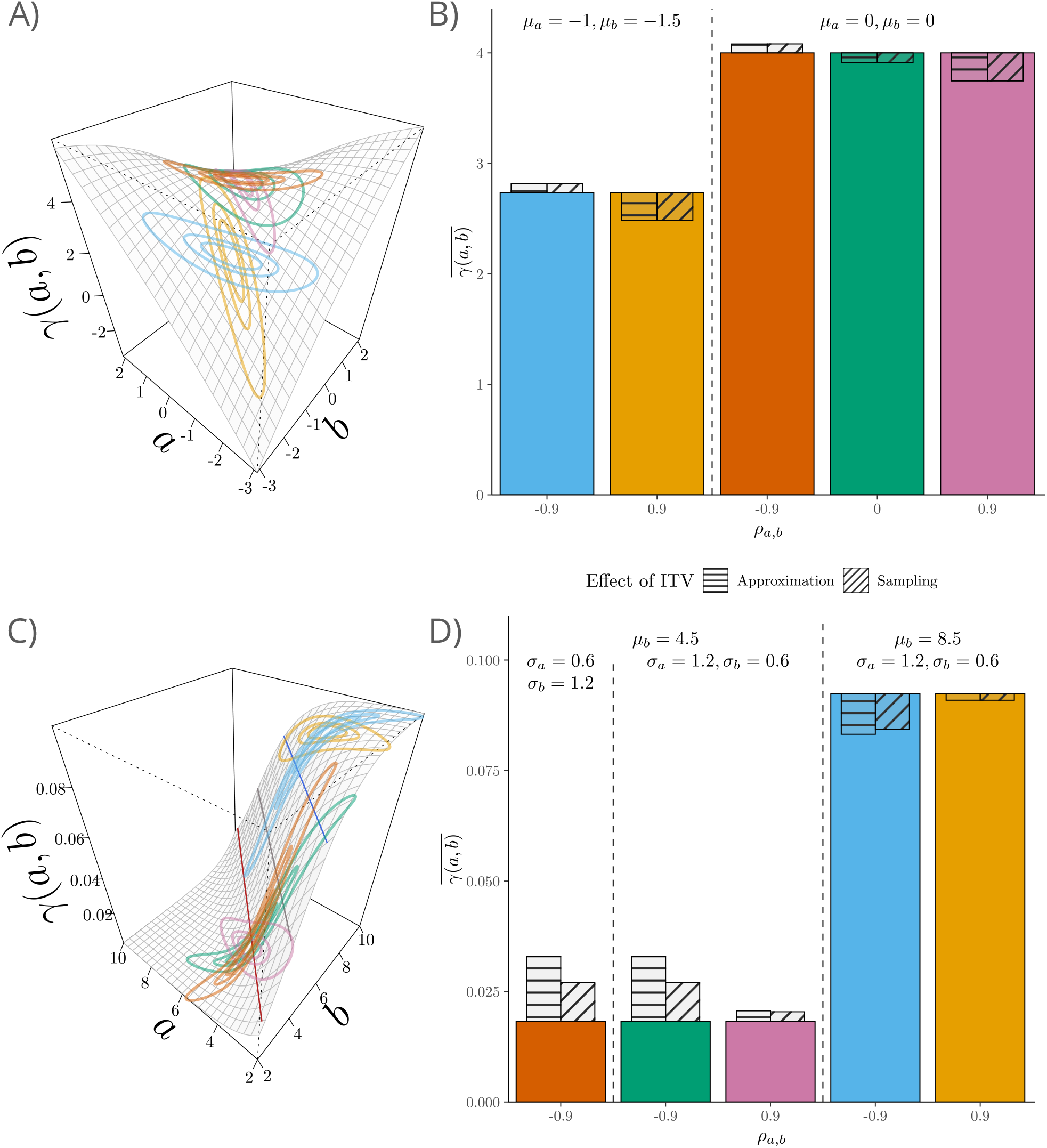
Effect of ITV on the average interaction parameter 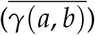 for the two interaction functions in Table 1. Left column: illustrations of the interaction functions *γ*. For the logistic (panel C) interaction function, lines of zero (grey), maximal (dark red) and minimal curvature (blue) are shown. Right column: the colored bars give the average interaction parameter without ITV or equivalently the naive estimate of the average interaction parameter *γ*(*µ*_*a*_, *µ*_*b*_). The striped and hatched bars represent how ITV changes the average interaction parameter according to the Taylor approximation and sampling approach compared to the naive case/the case without ITV. The joint trait distributions used in B and D are shown in A and C as three ellipses that contain 25%, 50% and 90% of all interactions, respectively, from inner to outer ellipse. (A–B) Polynomial with *p*_1_ = − 0.1, *p*_2_ = − 0.15, *p*_3_ = − 0.55, *p*_4_ = *p*_5_ = 0, *p*_6_ = 4, *σ*_*a*_ = 0.75, and *σ*_*b*_ = 0.5. (C–D) Logistic with *p*_max_ = 0.1, *k* = 1, *h* = 1.5 at *µ*_*a*_ = 4.5.

The second interaction function we explore is a *logistic interaction function* (figure 3C),

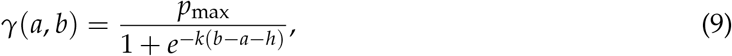

where *p*_max_ is the maximum interaction rate, *h* is the difference in traits at which the interaction rate is half the maximum and *k* determines the steepness of the curve. We chose this interaction function because it is non-negative everywhere, and it is commonly used for example to model various interactions where the rate or outcome depends on the difference in trait value between interacting individuals. For the sake of concreteness, we will assume in the following that the function represents the probability of a predator eating a prey when predation success increases with increasing predator size *b* relative to prey size *a*.

When the average predator is smaller than average prey size plus *h*, overall hunting efficiency is low, but trait variation in either species may yield occasional encounters between relatively large predators and relatively small prey, resulting in an increase in average hunting success (left three bars in figure 3D). This can be understood directly from the positive curvature with respect to both trait values in this region (figure 3C). The largest positive effect of ITV on the approximation occurs when the curvatures are maximal, on the line 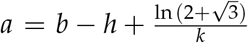 (dark red line in figure 3C, S6.2). The effect can be further enhanced with a negative correlation (left two bars in figure 3D). In this case, small predators are matched to large prey and obtain basically no food, but this is more than compensated for by large predators being matched to small prey that they can feed on efficiently. The effect of ITV on the average interaction parameter is reversed in the region where the average predator is more than *h* larger than the average prey (right two bars in figure 3D), with a maximum effect size in the approximation when 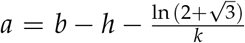 (blue line in figure 3C, S6.2). The effects of ITV vanish at the boundary between these regions (*b* − *a* = *h*, grey line in figure 3C), where the curvatures are zero, or when the difference between the average trait values becomes very large.

The logistic interaction function can be written purely in terms of the difference between trait values *a* and *b* and thus a change in variance in either species adds the same amount of variance to this difference. For such interaction functions, the approximation (Eq. (6)) can be simplified (see section S5):

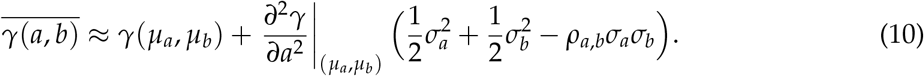

Hence, variation in *a* and *b* contributes equally to the difference between 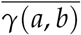 and the naive mean *γ*(*µ*_*a*_, *µ*_*b*_). The difference is enhanced if the traits are highly and negatively correlated (figure 3D), while the effect of ITV on the interaction function cancels out if the traits are perfectly correlated (correlation coefficient *ρ*_*a,b*_ = 1) and the trait variances are the same. Note that for this class of interaction functions the correlation cannot change the sign of the effect of ITV, which thus depends solely on 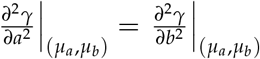 (section S5). For cases where the interaction function only depends on the difference in trait value, we thus reach the important general conclusion that, depending on the trait correlation, only taking into account variation in one species could underestimate the effect of ITV by a factor of four (with *σ*_*a*_ = *σ*_*b*_ = *σ* and *ρ*_*a,b*_ = −1, the effect of ITV would be estimated as *σ*^2^/2 when in fact it is 2*σ*^2^), but also falsely predict a nonzero effect of ITV that disappears when both species are studied (with *σ*_*a*_ = *σ*_*b*_ = *σ* and *ρ*_*a,b*_ = 1, the effect would be estimated as *σ*^2^/2 when in reality it is zero).

While in the polynomial interaction function, the approximation perfectly matches the exact estimate from the sampling approach, for the logistic interaction function, there is a discrepancy between approximation and sampling approach, but the approximation always correctly predicted the direction of the effect of ITV. As shown in figure S7.1, the Taylor approximation is close to the two exact approaches for small variances, but increasingly deviates for larger trait variances, as expected.

In the supporting information, we study two additional interaction functions. Firstly, the commonly used Gaussian interaction function that also depends only on the difference in trait values (S4). Secondly, we study a more complex and non-differentiable interaction function. This corresponds to a polynomial on which hard limits are imposed, so that it can never turn negative (section S3.2). While the sampling approach still works, the approximation does not perform well close to discontinuities in the curvature. For this interaction function, interestingly, the trait variation in species *B* affects the average interaction parameter through the correlation (last) term in the approximation (Eq. 6), even though the curvature of the interaction function with respect to *b* is zero.

### 2.4 Consequences for population dynamics and coexistence

Although the effects of ITV and trait correlations on average interaction parameters are already important on their own, in the final part of our framework we go one step further and explore how the resulting differences in average interaction parameters affect population dynamics and species coexistence (figure 1, right). The population models we use here all assume that the number of individuals in both species is large so that demographic stochasticity does not play a role and the cumulative effects of the random individual-by-individual trait-dependent interactions can be well captured by the average interaction parameter 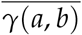. As a result, the model is de-terministic. In general, our approach allows one or more parameters of a dynamical system (in continuous or discrete time, though here we focus on continuous-time models) to be replaced by the average over trait-dependent interaction functions. We will now illustrate this application for a predator-prey model. A second detailed application, for a competition model, is given in section S10.

## 3 Application example 1: Qualitative and quantitative effects of ITV and interspecific trait correlations in a predator-prey model

Consider a predator-prey model:

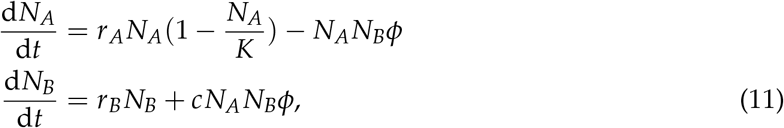

where *N*_*A*_ and *N*_*B*_ are the prey and predator abundances, *r*_*A*_ *>* 0 and *r*_*B*_ *<* 0 are the prey and predator intrinsic growth rates, *K* is the environmental carrying capacity, *c* is the conversion efficiency, and *φ* determines the interaction strength, which we assume to be trait-dependent, and thus set it to 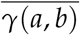. In this example, we will use the logistic interaction function (Eq. (9)) with the same parameters as in figure 3C. Since the interaction outcome for this function just depends on the difference between predator and prey trait values, it can also be displayed as in figure 4A.

**Figure 4.**
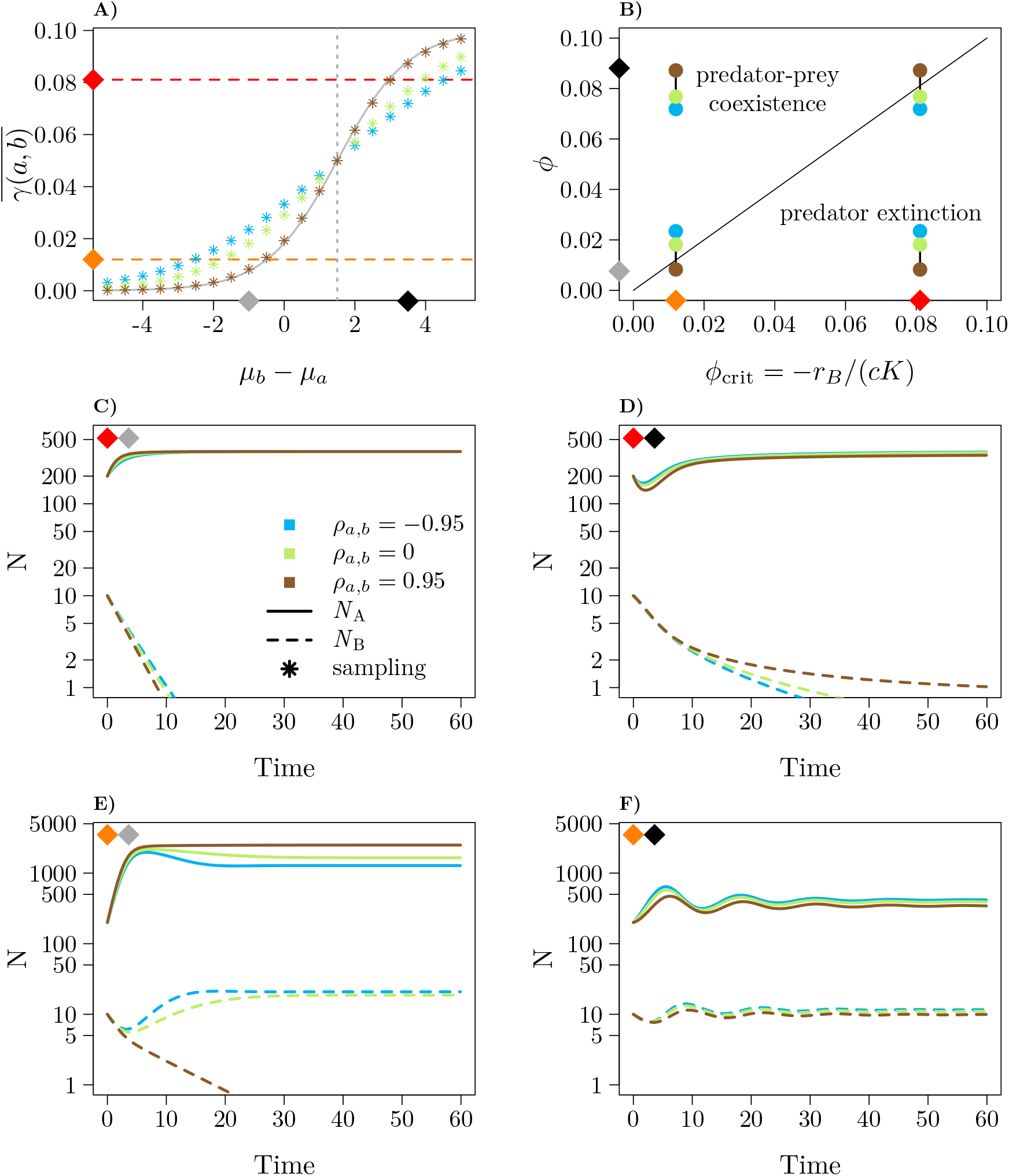
Effects of intraspecific trait variation and interspecific trait correlation on the attack rate *φ* and consequently on the dynamics and qualitative outcome of the predator-prey model (Eq. 11). Here *σ*_*a*_ = *σ*_*b*_ = 1.5 and the interaction is defined by a logistic interaction function with *p*_*max*_ = 0.1, *k* = 1, *h* = 1.5. The dotted vertical line is where the second derivative and thus the effect of ITV switches from positive to negative. The diamonds refer to different parameter settings: *K* = 370 and correspondingly a high value of *φ*_crit_ (red diamond), *K* = 2500 and correspondingly a low value of *φ*_crit_ (orange diamond), *µ*_*b*_ smaller than *µ*_*a*_ (grey diamond) and *µ*_*a*_ smaller than *µ*_*b*_ (black diamond). A) Estimates of the average interaction function at three values for the interspecific trait correlation using the sampling method. The grey line corresponds to the naive expectation and the case without ITV. B) Regions in parameter space with qualitatively different outcomes. C–F) Time series corresponding to four scenarios with the average attack rate as given by the sampling method. We do not show the lines corresponding to the case without ITV because they would overlap with the brown lines for *ρ*_*a,b*_ = 0.95. Other parameter values: *r*_*A*_ = 1, *r*_*B*_ = −0.3, *c* = 0.01.

The qualitative and quantitative effects of ITV and trait correlations on population dynamics and coexistence are shown in figure 4. As we have seen above (see Fig. 3 C,D), the effect of ITV in either species can be positive or negative depending on the difference in mean trait values (figure 4A). This effect can be almost eliminated by a strong positive correlation (thus it is no coincidence that the points for high *ρ*_*a,b*_ almost overlap the line without ITV) or amplified by a strong negative correlation. Even though the value of *ρ*_*a,b*_ cannot change the direction of the effect of ITV on the average interaction parameter for interaction functions of this form, it can still qualitatively affect the population dynamics. Coexistence in the model depends on whether the average attack rate parameter 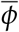 is above a critical threshold (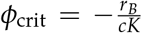, indicated by the dashed horizontal lines in figure 4A), which thus depends on the carrying capacity of the prey (S8). We show time series for high and low prey carrying capacity (red and orange diamonds), and for cases where either the prey mean trait *µ*_*a*_ is smaller than the predator mean trait *µ*_*b*_ (black diamond), or the other way around (grey diamond). In the scenario with the grey diamond, the difference in mean trait values *µ*_*b*_ − *µ*_*a*_ is below *h* (grey dashed line), ITV increases *φ* and this effect is enhanced by a strong negative trait correlation. At high carrying capacity, the minimum average interaction parameter that leads to coexistence is low (orange diamond) and ITV combined with a negative trait correlation can increase *φ* to a value above this low critical threshold (*φ*_crit_) and thereby promote coexistence, while without ITV or with a positive trait correlation the predators may go extinct (figure 4A,B,E). If, on the other hand, the predator-prey size difference is above *h* (black diamond), ITV reduces the interaction coefficient, an effect that is again stronger when combined with a strong negative trait correlation. If the coexistence threshold is relatively high (red diamond), this can drive the predator to extinction even if without ITV both species could coexist, unless a strong positive trait correlation reduces the effect of ITV (figure 4A,B,D). The potential for interspecific trait correlation to affect species coexistence is the largest when the general effect of ITV on the average interaction parameter is maximal (on the lines 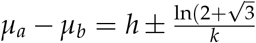, as explained above). Even if the change in average interaction parameter is not large enough to change the qualitative outcome, extinction times (red-grey diamonds) or equilibrium population abundances (orange-black diamonds) can be affected, although in this case the effect is only small (figure 4A,B,C,F, see S8.1 for more detailed information).

## 4 Different parametrization options

Before moving on to the next application example, we would like to point out one subtlety about our approach: There is not always just one correct way of setting up or estimating the interaction function and joint trait distribution, but there can be multiple equivalent ways that can be more or less convenient depending on the research question and study design. In fact, it is always possible to account for all the information about occurrence or encounter patterns in the interaction function and work with the marginal distributions, i.e. have no correlation, since the average interaction parameter can be rewritten as

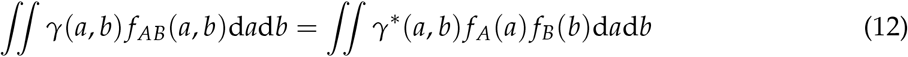

with a new interaction function

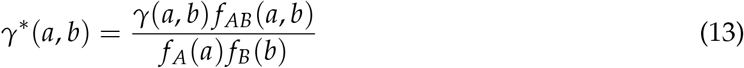

and marginal trait distributions *f*_*A*_(*a*) and *f*_*B*_(*b*).

Thus, the joint trait distribution and the interaction function allow us to parameterize the system in different ways depending on what is easiest and most meaningful given the biological system and potential empirical data. We will illustrate these different parametrization options for a predator-prey system. A first way to parameterize the system is to assume that the trait values of interacting predator and prey individuals are independently drawn from their respective trait distributions, i.e. the joint trait distribution is the product of the two marginal distributions (*f*_*A*_(*a*) and *f*_*B*_(*b*)). The interaction function then gives the rate and outcome of an interaction of the randomly generated pair of individuals given their trait values. This parametrization corresponds to the right hand side of Eq. (12) and makes sense if interactions between individuals indeed occur randomly with respect to traits. If individuals interact non-randomly with respect to traits, for example because predators and prey with larger trait values tend to co-occur more often in the same micro-environments or encounter each other more often, it can still be used, but the non-random interaction then needs to be modeled as part of the interaction function. In the predatorprey example, the interaction function then can be understood as integrating the probability that individuals with certain trait values co-occur, the rate at which they encounter each other, and the probability that the predator successfully kills the prey. Such a parametrization can make sense if one has data on the marginal trait distribution, but no data on co-occurrence and encounter patterns.

If one does have such data available, it can make more sense to parameterize the system using a joint trait distribution function with non-zero correlation (left hand side of Eq. (12)). For example, if one has data on trait-dependent spatial or temporal co-occurrence patterns, the joint trait distribution would specify how likely the pairs of individuals with different trait combinations are to co-occur, and the interaction function would model the rate at which co-occurring individuals encounter each other and the predator then successfully captures the prey. If one even has data on trait patterns of pairs of individuals that encounter each other, then the joint trait distribution could be parameterized as the distribution of pairs of trait values in predator-prey encounters and the interaction function as the overall encounter rate times the probability that the predator successfully kills the prey given an encounter. But we have to be cautious because the joint trait distribution will have the trait distributions of the individual species as marginal distributions only if the probability that an individual is part of some interaction pair does not depend on traits. While parts of our framework still work if this assumption is not met, others like predicting interaction outcomes for different trait variances will not. So, in this case we recommend to also obtain the marginal trait distributions for the individual species and work with uncorrelated trait distributions.

## 5 Application example 2: Estimating the effects of ITV from field data

Here we use simulated field data to illustrate how the joint trait distribution, interaction function and consequentially the effects of intraspecific trait variation can be estimated from empirical data. Consider a hypothetical field site structured into 100 micro-habitats and inhabited by two interacting populations: a predator population and a prey population. To generate our artificial data set, we assumed the mechanism outlined in section 2.1.1 where individuals exhibit phenotypic plasticity in response to an environmental factor in their micro-environment (Eq. 1). We assumed that the values of this environmental factor are drawn from a normal distribution with mean 0 and standard deviation 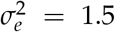. In each micro-habitat, there are 400 prey individuals and 10 predator individuals. The random individual trait effect has standard deviation *σ*_*ϵ,a*_ = *σ*_*ϵ,b*_ = 0.7 in both species. We considered two scenarios, one in which *h*_*A*_ = 4, *h*_*B*_ = 4, *m*_*A*_ = 0.5, *m*_*B*_ = −0.6, i.e. prey and predator respond in opposite ways to the micro-environment, which leads to a negative correlation among prey and predator trait values in interacting individuals (figure 5A), and one where prey and predator respond similarly with *h*_*A*_ = 2, *h*_*B*_ = 4, *m*_*A*_ = 0.5, *m*_*B*_ = 0.6, which leads to a positive correlation (figure 5B). We assume that the encounter rate for each pair of individuals in the same patch is 0.001. With an observation time of 1 time unit, the number of encounters for each pair of individuals in the same patch is then Poisson-distributed with mean 0.001. Given an encounter, the predator kills the prey with a probability based on the logistic interaction function (with *k* = 1 and *h* = 1.5 like in figure 4 A, but with *p*_*max*_ = 1). Our pseudo-observed data set then contains all the encounters and for each encounter the prey individual’s trait value, the predator individual’s trait value, and whether or not the predator was successful in killing the prey. Note that we assume that prey individuals that are killed are replaced by another individual with the same trait value such that the population size and the trait distributions stay constant over time.

**Figure 5.**
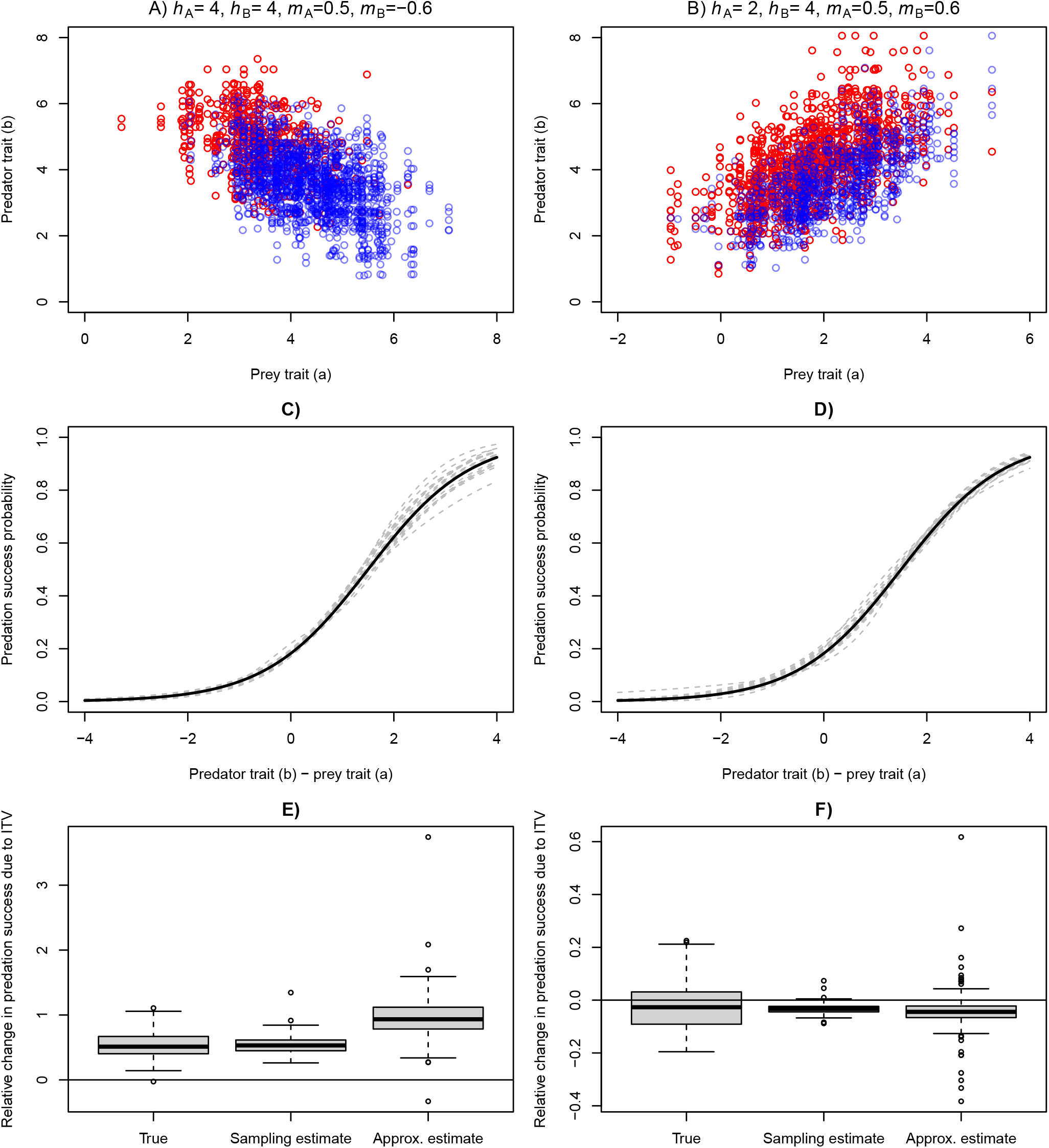
Illustration of our approach of estimating the impact of ITV on average predation success from field data. The two columns show two scenarios that differ in the reaction norms via which individuals respond to their micro-environment. A, B) Trait values of pairs of interacting individuals in the first replicate, with successful kills shown in red and encounters where the prey escaped shown in blue. Across all replicates in A, there were on average 2191.5 encounters, with 610.8 kills and 1580.7 escapes. In B, there were on average 2187.7 encounters, with 1317.7 kills and 870.1 escapes. C, D) Estimated interaction function for the first 20 replicates shown in dashed gray lines. Note that we have evaluated the two-dimensional interaction function at the respective mean prey trait *h*_*A*_. The true interaction function is shown in black. E, F) Boxplots of true and estimated effects of ITV on total predation success among the 100 replicates.

Given such a pseudo-observed data set, our final goal is to estimate the impact of ITV on total predation success, i.e., the total number of kills made, of course assuming that the true input parameters are not known. For this, we first fit an interaction function 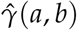 by fitting a tensor product smooth to the data using the `mgcv` library (Wood, 2006) in R (R Core Team, 2022). The tensor product smooth does not make any prior assumptions about the shape of the interaction function and allows for interactive effects. In the fit, we use family “binomial” and a logit link because our predation success data is binary. We then randomly sample 500, 000 new pairs of trait values for interacting individuals from a multivariate normal distribution parameterized with the means, standard deviations and the correlation coefficient estimated directly from the observed trait values across all the encounters. By then averaging the 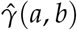 values of the sampled trait-value pairs, we get a sampling estimate of the average interaction parameter.

To obtain a Taylor approximation, we use a numerical method (the `hessian` function from the `numDeriv` package) to obtain the second partial derivatives from the estimated interaction function 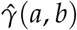. We then plug these curvatures and the estimated distribution parameters into (6). To obtain the estimated mean predation rate without ITV, we simply plug the estimated mean trait values 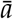 and 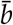 into the estimated interaction function to obtain 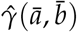. The estimated effect of ITV relative to the case without ITV is then the average predation success with ITV minus the average predation success without ITV, 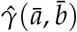, divided by 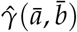. If this were a real field study, we would be done now and state this result or we might do some bootstrapping, but there is usually no way to verify it in the field. With simulated data, however, we can check how well we have done by also simulating a second data set without ITV, where all individuals of species *i* have a trait equal to the mean trait as given by Eq. (2), and analogously quantifying the relative change in predation success due to ITV. Since there is a lot of stochasticity in the true data, we ran this whole experiment 100 times. That is, for each replicate, we had one simulation run without ITV and one with ITV. While the estimates of the effect of ITV were only based on the simulation run with ITV, the “true” value of the effect of ITV was based on comparing these two runs.

Across the 100 replicates, the shape of the interaction function is generally estimated very well (figure 5 C, D) and the median relative change in predation success due to ITV is well approximated by the sampling method (figure 5 E,F). In the first scenario, it correctly predicts a strong positive effect of ITV on predation success, in the second scenario a weaker negative effect. In the first scenario, the approximation somewhat overestimates the effect of ITV, while in the second scenario it captures the effect of ITV well. The results are qualitatively similar in cases with lower or no environmental variance *σ*_*e*_ (figure S9.1 and S9.2), but since a lower *σ*_*e*_ leads to lower ITV and weaker correlations, the effects of ITV are overall weaker. This experiment shows that, given sufficient data, it is possible in principle to estimate both the shape of the interaction function and the impact of ITV on average interacting parameters just from a data set of pairs of interacting individuals and the outcome of each interaction.

## 6 Discussion

### 6.1 General modeling framework

Our modeling approach has two key ingredients, the joint trait distribution and the interaction function, based on which the impacts of ITV and trait correlations can be estimated in different ways. Exact estimates can be obtained from a random sampling method or other types of numerical integration over the interaction function with respect to the joint trait distribution. On the other hand, the Taylor approximation is computationally more efficient, depends only on local estimates of the curvatures, and provides additional insight into how the different components of ITV contribute to the average interaction parameter. While the approximation is exact when curvatures are constant, as is the case for a second-order polynomial, it may be inaccurate when the curvatures change significantly over the part of trait space where the bulk of the interactions take place. For example in our non-differentiable interaction function, the approximation could even predict an effect in the opposite direction whenever interactions took place on both sides of a discontinuity (see section S3.2). For the logistic and Gaussian interaction functions, the secondorder Taylor approximation always correctly predicted the direction of the change compared to the naive value where the species’ mean trait values are plugged into the interaction function or equivalently to the case without ITV (figure 3 and figure S4.1). However, when variances were large, the approximation tended to overestimate their impact, potentially leading to wrong con-clusions regarding species coexistence. Note that higher order terms can be straightforwardly included into the approximation when traits are bivariate normal (as shown in S2.3). The higher order terms improve the accuracy when trait variances are small, but they may decrease the accuracy when trait variances are large (figure S2.1). This behaviour is expected because, away from the point at which the approximation is performed, Taylor approximations do not necessarily converge with increasing number of terms (see for example §6.2.1 in Witelski and Bowen, 2015).

Taylor approximations have been used to link trait variation to population dynamics before, both in single-species models (Bjørnstad and Hansen, 1994) and in models with two competing species with an underlying trait, that affects inter- and intraspecific competition simultaneously (Hart et al., 2016). More generally, such Taylor approximations have also been used to study the effect of spatio-temporal variation in environmental conditions on species’ growth rates, as well as population and community dynamics (e.g. Bernhardt et al. 2018; Denny 2019; Gravel et al. 2011; Koussoroplis et al. 2017; and the “scale-transition theory” in modern coexistence theory Barabás et al. 2018; Chesson 2012; Chesson et al. 2005; Melbourne and Chesson 2005). For example, in modern coexistence theory, the effect of spatial or temporal environmental heterogeneity on fitness is captured well by the product of local curvature of the fitness function and environmental variance as long as the environmental variability is small relative to the scale over which the curvature of the fitness function changes. In other cases, other properties of the distribution (e.g. frequency of extreme environmental conditions) can become important and one would need full knowledge of the distribution function and fitness function to quantify the effects. Our Taylor approximation shares the strengths and weaknesses of these earlier approaches in that it also provides intuitive results, in our case for the joint effects of trait variation and correlations in two interacting species, but depends on accurate estimates of the curvatures and trait variation being small relative to the scale of the interaction function.

### 6.2 The effect of ITV and trait correlations on population dynamics and coexistence

Our results show how ITV in each of two interacting species can affect average interaction parameters such as predation rates through nonlinear averaging over the joint trait distribution. The magnitude and direction of this effect depend on the shape of the interaction function, average trait values, amounts of trait variation in each species, as well as on the trait correlation between interacting individuals. As we have seen in our first application example, these nonlinear averaging effects can change the joint population dynamics both quantitatively (e.g. influencing equilibrium population densities) and qualitatively (species coexistence or extinction).

One class of interaction functions we have looked at in detail are those for which interaction outcomes only depend on the difference in trait values. This seems to be roughly the case for size-based predator-prey interaction when size is measured on a logarithmic scale (Portalier et al., 2019). We have shown that for such interaction functions trait correlations can nullify or double the effect of ITV on the average interaction function. This can lead to qualitative differences in the resulting population dynamics. For example in our predator-prey model, strong positive trait correlations could prevent or cause predator extinctions, depending on the difference in mean trait value between prey and predator. When predators are sufficiently larger or faster than prey (black diamond in figure 4) and thus, on average, very efficient at capturing them, a positive correlation ensures that each predator is matched to a prey that it can hunt effectively. Such a correlation could emerge from co-occurrence patterns or via behavioral choices of the predators to only pursue prey items that are not too large but also worth the energy expenditure of hunting, capturing, and handling them (Portalier et al., 2019). In particular when prey have a small carrying capacity or when predators need a large number of prey items to survive, such positive correlation can prevent the predators from going extinct. Conversely when prey are relatively large or fast relative to predators such that predators are generally poor at consuming prey, while at the same time the critical predation rate is low because prey are abundant or pro-vide a lot of energy per capture, a negative correlation ensures that at least the large predators are matched to a prey that they can hunt effectively, thereby preventing the predator from going extinct (figure 4). If we had instead used a Gaussian interaction function, the average attack rate parameter would increase with increasing trait correlations when trait means are close together, but decrease when trait means are far apart, regardless of the sign of the difference. Such a mismatch-based interaction function has been used for example in a Rosenzweig-MacArthur model where an increase in variation in attack rate and handling time tends to promote coexistence when trait mismatch is large and hinder coexistence when trait mismatch is small (Gibert and Brassil, 2014; Gibert and DeLong, 2015).

In an additional example in SI S10, we similarly explored the effects of ITV and trait correlations in a competition model. This example also illustrates that multiple model parameters can be simultaneously affected by ITV and trait correlations, in this case the interspecific competition coefficient and the two intraspecific competition coefficients. While many of the results recapitulate earlier findings by Hart et al. (2016) and Barabás and D’Andrea (2016), we also made a few new observations. For example, when the trait means of the competing species are close together, ITV in either species can promote coexistence in some cases where interspecific competition width is smaller than intraspecific competition width (see figure S10.3).

### 6.3 Estimating the effects of ITV from data

So far, there have been studies examining the impact of ITV in a single species on average interaction parameters from laboratory experimental data or a mix of field data and laboratory data (Coblentz et al., 2021; Pearse et al., 2018; Wetzel et al., 2016). Furthermore, (Hausch et al., 2018) performed an experiment where they investigated the consequences of ITV in two competing species on species invasion growth rates and coexistence. However, to our knowledge, it is so far not clear how the effect of joint ITV in two interacting species can be estimated based on field data. Being able to do this would help assess the importance of ITV in natural ecological communities and thus for biodiversity conservation.

In our second application example, we have thus used simulated data to illustrate how the impact of ITV in two interacting species on average interaction parameters can be estimated from field data. In this example, the available data set consisted of observations of encounter events between predator individuals and prey individuals. For each observed encounter, we had the trait value of the prey individual, the trait value of the predator and the outcome of the interaction, i.e., whether the predator successfully killed the prey. In this case, it was natural to parameterize the joint trait distribution as the distribution of pairs of trait values in individuals that encounter each other and to parameterize the interaction function as the outcome of the predation attempt given that individuals with certain trait values encounter each other.

However, in other cases one might only have data on marginal trait distributions of each species and trait values of both individuals for successful predation events, e.g. from predator gut content analysis. In such cases, one can assume that both trait values are drawn independently (i.e. the joint trait distribution is the product of the marginal distributions) and the interaction function then describes both the probability that two individuals with given trait values actually interact, for example the probability that a predator with trait value *a* encounters a prey individual with trait value *b* and the outcome of the interaction.

The benefit of this flexibility is that the model can be parameterized in different ways depending on what is most convenient given the experimental design or available field data. If sufficient data are available, we recommend parametrizing the model with a joint trait distribution with potential trait correlations as this can give more mechanistic insights into the effects of ITV compared to a model where trait effects on co-occurrence, encounter, and outcomes of the encounter are all lumped together into the same interaction function.

If one assumes a bivariate normal distribution, the parameters of the joint trait distribution can be obtained simply by computing means, variances and covariance from the observed trait data. Other distributions can be fit in similarly if necessary. However, one could also work with a (smoothed) empirical distribution to avoid assumptions on the shape of the joint trait distribution. To fit the interaction function, we recommend using a flexible approach such as a tensor product smooth (Wood, 2006) that does not make strong *a priori* assumptions on the shape of the interaction function. Although this approach is more data-demanding than approaches where a certain functional form is assumed (as e.g. in Bartomeus et al., 2016), it has the advantage that the curvature of the interaction function, which ultimately determines the direction of the effect of ITV, emerges from the data, thereby avoiding bias in a certain direction.

While applying our approach to real field data is beyond the scope of this article, there are already some promising data sets containing individual observations of interaction outcomes and the trait values of the interacting partners. For example, pollination success has been recorded for plants with different nectar-holder morphology interacting with pollinators with various beak or proboscis lengths (Ibanez, 2012; Missagia and Alves, 2018). Similarly, data are available on trait matching between predator feeding and prey vulnerability (Brousseau et al., 2018), insects and deadwood’s chemical compound traits in detritivore networks (Wende et al., 2017), and morphological plant and animal traits in plant-frugivore (Bender et al., 2018) or detritivore (Brousseau et al., 2019) networks. Bartomeus et al. (2016) have indeed fitted Gaussian interaction functions on datasets describing fish predator-prey body size (Barnes et al., 2008), grasshopper incisive strength and leaf dry matter relationship (Deraison et al., 2015), and plant-pollinator data (Bartomeus et al., 2008). While many of these examples pool multiple species within a functional group, our approach should still be usable. Despite these examples, more such data on individual by individual interactions are needed in order to better understand the effect of ITV on species interactions.

### 6.4 Outlook

There are multiple ways in which our approach can be extended to include more biological realism. First, our approach could be straightforwardly extended to allow for multiple traits affecting different ODE parameters or systems with more than two species.

Second, like many previous studies (e.g. Begon and Wall, 1987; Stump et al., 2022), we have assumed static non-heritable trait distributions. These can arise through stable age or stage struc-tures (as expected in matrix models and integral projection models, see for example Caswell, 2001; Easterling et al., 2000) or through phenotypic plasticity in response to a distribution of microenvironments, as explained in the derivation above (Moran et al., 2016). In the latter case we would expect the trait correlation patterns also to remain constant over time. However, one might expect the joint trait distribution to change both directly as well as over multiple generations, for example if prey with certain trait values are being consumed more frequently, or individuals with a certain trait value face stronger competition than others, or simply because of variable environmental conditions driving plastic trait variation (Jackson and Xue, 2024). To allow for changing trait distributions over time, we could combine our framework with one of the existing approaches for modeling changes in trait distributions. To deal for example with the gradual loss of prey items in a predator-prey setting, we could employ approaches similar to those developed for estimating functional response curves from experimental data (DeLong, 2021). Assuming asexual reproduction, we could use partial differential equations or moment equations to model the eco-evolutionary dynamics of population sizes and trait distributions (see e.g. Senthilnathan and Gavrilets, 2021; Wickman et al., 2023). For systems where heritability is not perfect, but the prey can be described in terms of two genetic types with different trait means and remaining plastic trait variation in each type (see e.g. Okuyama, 2011; Schreiber et al., 2011), we can average the interaction between each of the prey types and the predator and still obtain an intuitive understanding of the effect of ITV on the average interaction parameters. Alternatively, we could allow for temporally changing trait values in either species by calculating a selection gradient from the interaction function and letting the mean trait value evolve accordingly (e.g. Barabás and D’Andrea, 2016; Bengfort et al., 2017; Lande, 1976). At any given time step we can then calculate the effect of ITV on the average interaction function, based on the mean trait values, and we can use the approximation to speed up the process. Models that allow for dynamic shifts in trait distributions can sometimes exhibit quite different behavior from those with fixed trait distribution (Jackson and Xue, 2023).

Third, another assumption of the framework is that the total interaction parameter can be written as the sum of independent individual-by-individual interactions. However, in some important ecological models, this is not the case. For example in predator-prey systems with predator saturation (such as models with a type II functional response), whether or not a predator individual eats a particular prey individual can depend on how many prey individuals it has consumed previously and thus also on their traits. While ITV just in the predator can be incorporated relatively easily (see e.g. Gibert and Brassil, 2014; Okuyama, 2008; Schreiber et al., 2011), extending our approach to including trait variation in both species and interspecific trait correlation into such models requires further research. Alternatively to nonlinear averaging, one could also explicitly keep track of the number of hunting and the number of handling predators separately to account for predator saturation (Okuyama, 2012). Non-independence between pairwise interactions may also occur. For example, in plant communities, associational effects are often observed where the herbivory pressure on a focal plant depends on its neighbors and their traits, e.g. their level of defense or ontogenetic stage (Cope et al., 2020; Hahn and Orrock, 2016). Furthermore, individuals may adjust their physiology or behavior in response to an interaction partner, which then affects the outcome of subsequent interactions with other individuals (e.g. physiological tracking of herbivores, Pearse et al., 2018). Developing a general framework that describes such non-additive cases coherently is an important direction for future research.

### 6.5 Conclusion

Our approach can be applied to a large variety of interaction functions and joint trait distributions. It allows for a straightforward and rapid assessment of the expected effect of variation in two species and interspecific trait correlations on the average interaction parameter. We have furthermore shown that such effects can strongly affect population dynamics in the example of a predator-prey system and can even make or break species coexistence. Finally, we have shown that by estimating the joint trait distribution and the interaction function and then applying our approach, the impact of ITV can in principle be estimated from field data.

## Acknowledgements

This work was supported by the German Research foundation DFG (project numbers 316099922 and 396782288 within SFB TRR 212 and WI 4544/2-1 within FOR 3000).

## Data availability

The R scripts to generate the results are supplied as supporting information. No external data were used.

## Supporting information

**S1–S10** contain additional analyses and results. All **R code** used for the analyses is available at https://doi.org/10.6084/m9.figshare.27900639.v1.

## Literature Cited

Barabàs, G., and R. D’Andrea. 2016. The effect of intraspecific variation and heritability on community pattern and robustness. Ecology Letters 19:977–986.

Barabàs, G., R. D’Andrea, and S. M. Stump. 2018. Chesson’s coexistence theory. Ecological Monographs 88:277–303.

Barnes, C., D. Bethea, R. Brodeur, J. Spitz, V. Ridoux, C. Pusineri, B. Chase, M. Hunsicker, F. Juanes, A. Kellermann, et al. 2008. Predator and prey body sizes in marine food webs: Ecological Archives e089-051. Ecology 89:881–881.

Bartomeus, I., D. Gravel, J. M. Tylianakis, M. A. Aizen, I. A. Dickie, and M. Bernard-Verdier. 2016. A common framework for identifying linkage rules across different types of interactions. Functional Ecology 30:1894–1903.

Bartomeus, I., M. Vilà, and L. Santamaría. 2008. Contrasting effects of invasive plants in plant– pollinator networks. Oecologia 155:761–770.

Begon, M., and R. Wall. 1987. Individual variation and competitor coexistence: a model. Functional Ecology 1:237–241.

Bender, I. M., W. D. Kissling, P. G. Blendinger, K. Böhning-Gaese, I. Hensen, I. Kühn, M. C. Muñoz, E. L. Neuschulz, L. Nowak, M. Quitiàn, et al. 2018. Morphological trait matching shapes plant–frugivore networks across the Andes. Ecography 41:1910–1919.

Bengfort, M., E. van Velzen, and U. Gaedke. 2017. Slight phenotypic variation in predators and prey causes complex predator-prey oscillations. Ecological Complexity 31:115–124.

Bernhardt, J. R., J. M. Sunday, P. L. Thompson, and M.I. O’Connor. 2018. Nonlinear averaging of thermal experience predicts population growth rates in a thermally variable environment. Proceedings of the Royal Society B: Biological Sciences 285:20181076.

Bjørnstad, O. N., and T. F. Hansen. 1994. Individual variation and population dynamics. Oikos 69:167–171.

Bolnick, D. I., P. Amarasekare, M. S. Araújo, R. Bürger, J. M. Levine, M. Novak, V. H. Rudolf, S. J. Schreiber, M. C. Urban, and D. A. Vasseur. 2011. Why intraspecific trait variation matters in community ecology. Trends in Ecology & Evolution 26:183–192.

Brousseau, P.-M., D. Gravel, and I. T. Handa. 2018. Trait matching and phylogeny as predictors of predator–prey interactions involving ground beetles. Functional Ecology 32:192–202.

Brousseau, P.-M., D. Gravel, and I. T. Handa. 2019. Traits of litter-dwelling forest arthropod predators and detritivores covary spatially with traits of their resources. Ecology 100:e02815.

Caswell, H. 2001. Matrix Population Models: construction, analysis and interpretation. 2nd ed. Sinauer Associates, Inc. Publishers, Sunderland, MA, USA.

Chesson, P. 2012. Scale transition theory: Its aims, motivations and predictions. Ecological Complexity 10:52–68.

Chesson, P., M. J. Donahue, B. A. Melbourne, and A. L. W. Sears. 2005. Scale transition theory for understanding mechanisms in metacommunities. Pages 279–306 in M. Holyoak, M. A. Leibold, and R. D. Holt, eds. Metacommunities: Spatial Dynamics and Ecological Communities. The University of Chicago Press, Chicago, IL, USA.

Clausen, A., and S. Sokol. 2020. Deriv: R-based Symbolic Differentiation. Deriv package version 4.1.

Coblentz, K. E., S. Merhoff, and M. Novak. 2021. Quantifying the effects of intraspecific variation on predator feeding rates through nonlinear averaging. Functional Ecology 35:1560–1571.

Cope, O. L., Z. Becker, P. J. Ode, R. L. Paul, and I. S. Pearse. 2020. Associational effects of plant ontogeny on damage by a specialist insect herbivore. Oecologia 193:593–602.

DeLong, J. P. 2021. Predator Ecology: Evolutionary Ecology of the Functional Response. Oxford University Press Oxford.

Denny, M. 2019. Performance in a variable world: using Jensen’s inequality to scale up from individuals to populations. Conservation Physiology 7:coz053.

Deraison, H., I. Badenhausser, L. Börger, and N. Gross. 2015. Herbivore effect traits and their impact on plant community biomass: an experimental test using grasshoppers. Functional Ecology 29:650–661.

Des Roches, S., D. M. Post, N. E. Turley, J. K. Bailey, A. P. Hendry, M. T. Kinnison, J. A. Schweitzer, and E. P. Palkovacs. 2018. The ecological importance of intraspecific variation. Nature Ecology & Evolution 2:57–64.

Easterling, M. R., S. P. Ellner, and P. M. Dixon. 2000. Size-specific sensitivity: applying a new structured population model. Ecology 81:694–708.

Fridley, J. D., J. P. Grime, and M. Bilton. 2007. Genetic identity of interspecific neighbours mediates plant responses to competition and environmental variation in a species-rich grassland. Journal of Ecology 95:908–915.

Genung, M. A., J. K. Bailey, and J. A. Schweitzer. 2012. Welcome to the neighbourhood: interspecific genotype by genotype interactions in Solidago influence above- and belowground biomass and associated communities. Ecology Letters 15:65–73.

Gibert, J. P., and C. E. Brassil. 2014. Individual phenotypic variation reduces interaction strengths in a consumer-resource system. Ecology and Evolution 4:3703–3713.

Gibert, J. P., and J. P. DeLong. 2015. Individual variation decreases interference competition but increases species persistence. Pages 45–64 in S. Pawar, G. Woodward, and A. I. Dell, eds. Advances in Ecological Research: Trait-Based Ecology - From Structure to Function, vol. 52, 1st ed. Elsevier Ltd., Waltham, MA, USA.

Gonzàlez-Varo, J. P., and A. Traveset. 2016. The labile limits of forbidden interactions. Trends in Ecology &amp; Evolution 31:700–710.

Gravel, D., F. Guichard, and M. E. Hochberg. 2011. Species coexistence in a variable world. Ecology Letters 14:828–839.

Hahn, P. G., and J. L. Orrock. 2016. Neighbor palatability generates associational effects by altering herbivore foraging behavior. Ecology 97:2103–2111.

Halbritter, A. H., S. Fior, I. Keller, R. Billeter, P. J. Edwards, R. Holderegger, S. Karrenberg, A. R. Pluess, A. Widmer, and J. M. Alexander. 2018. Trait differentiation and adaptation of plants along elevation gradients. Journal of Evolutionary Biology 31:784–800.

Hart, S. P., S. J. Schreiber, and J. M. Levine. 2016. How variation between individuals affects species coexistence. Ecology Letters 19:825–838.

Hausch, S., S. M. Vamosi, and J. W. Fox. 2018. Effects of intraspecific phenotypic variation on species coexistence. Ecology 99:1453–1462.

Holdridge, E. M., and D. A. Vasseur. 2022. Intraspecific variation promotes coexistence under competition for essential resources. Theoretical Ecology 15:225–244.

Ibanez, S. 2012. Optimizing size thresholds in a plant–pollinator interaction web: towards a mechanistic understanding of ecological networks. Oecologia 170:233–242.

Inouye, B. D. 2005. The importance of the variance around the mean effect size of ecological processes: Comment. Ecology 86:262–265.

Jackson, Z., and B. Xue. 2023. Heterogeneity of interaction strengths and its consequences on ecological systems. Scientific Reports 13:1905.

Jackson, Z., and B. Xue. 2024. Dynamic trait distribution as a source for shifts in interaction strength and population density. The American Naturalist 204:1–14.

Jensen, J. L. W. V. 1905. Sur les fonctions convexes et les inègalitès entre les valeurs moyennes. Acta Mathematica 30:175–193.

Koussoroplis, A.-M., T. Klauschies, S. Pincebourde, D. Giron, and A. Wacker. 2019. A comment on “Variability in plant nutrients reduces insect herbivore performance”. Rethinking Ecology 4:79–87.

Koussoroplis, A.-M., S. Pincebourde, and A. Wacker. 2017. Understanding and predicting physiological performance of organisms in fluctuating and multifactorial environments. Ecological Monographs 87:178–197.

Lande, R. 1976. Natural selection and random genetic drift in phenotypic evolution. Evolution 30:314–334.

Maynard, D. S., C. A. Servàn, J. A. Capitàn, and S. Allesina. 2019. Phenotypic variability promotes diversity and stability in competitive communities. Ecology Letters 22:1776–1786.

McPeek, M. A., S. J. McPeek, and F. Fu. 2022. Character displacement when natural selection pushes in only one direction. Ecological Monographs 92:e1547.

Melbourne, B. A., and P. Chesson. 2005. Scaling up population dynamics: integrating theory and data. Oecologia 145:179–187.

Missagia, C. C. C., and M. A. Alves. 2018. Does beak size predict the pollination performance of hummingbirds at long and tubular flowers? A case study of a neotropical spiral ginger. Journal of Zoology 305:1–7.

Moran, E. V., F. Hartig, and D. M. Bell. 2016. Intraspecific trait variation across scales: implications for understanding global change responses. Global Change Biology 22:137–150.

Moran, N. P., B. A. Caspers, N. Chakarov, U. R. Ernst, C. Fricke, J. Kurtz, N. D. Lilie, L. K. Lo, C. Müller, R. R, E. Takola, P. C. Trimmer, K. J. van Benthem, J. Winternitz, and M. J. Wittmann. 2022. Shifts between cooperation and antagonism driven by individual variation: a systematic synthesis review. Oikos 2022:e08201.

Nakazawa, T. 2017. Individual interaction data are required in community ecology: a conceptual review of the predator–prey mass ratio and more. Ecological Research 32:5–12.

Okuyama, T. 2008. Individual behavioral variation in predator–prey models. Ecological Research 23:665–671.

Okuyama, T.. 2011. Individual variation in prey choice in a predator-prey community. Theoretical Population Biology 79:64–69.

Okuyama, T.. 2012. Behavioral states of predators stabilize predator-prey dynamics. Theoretical Ecology 5:605–610.

Pearse, I. S., R. Paul, and P. J. Ode. 2018. Variation in plant defense suppresses herbivore performance. Current Biology 28:1981–1986.e2.

Pettorelli, N., T. Coulson, S. M. Durant, and J.-M. Gaillard. 2011. Predation, individual variability and vertebrate population dynamics. Oecologia 167:305–314.

Pettorelli, N., A. Hilborn, C. Duncan, and S. M. Durant. 2015. Individual variability: The missing component to our understanding of predator-prey interactions. Pages 19–44 in Advances in Ecological Research: Trait-based ecology - From structure to function, vol. 52. Elsevier.

Portalier, S. M. J., G. F. Fussmann, M. Loreau, and M. Cherif. 2019. The mechanics of predator–prey interactions: First principles of physics predict predator–prey size ratios. Functional Ecology 33:323–334.

R Core Team. 2022. R: A Language and Environment for Statistical Computing. R Foundation for Statistical Computing, Vienna, Austria.

Raffard, A., F. Santoul, J. Cucherousset, and S. Blanchet. 2019. The community and ecosystem consequences of intraspecific diversity: a meta-analysis. Biological Reviews 94:648–661.

Ruel, J. J., and M. P. Ayres. 1999. Jensen’s inequality predicts effects of environmental variation. Trends in Ecology & Evolution 14:361–366.

Schirmer, A., J. Hoffmann, J. A. Eccard, and M. Dammhahn. 2020. My niche: individual spatial niche specialization affects within- and between-species interactions. Proceedings of the Royal Society B: Biological Sciences 287:20192211.

Schreiber, S. J., R. Bürger, and D. I. Bolnick. 2011. The community effects of phenotypic and genetic variation within a predator population. Ecology 92:1582–1593.

Senthilnathan, A., and S. Gavrilets. 2021. Ecological consequences of intraspecific variation in coevolutionary systems. The American Naturalist 197:1–17.

Stump, S. M., C. Song, S. Saavedra, J. M. Levine, and D. A. Vasseur. 2022. Synthesizing the effects of individual-level variation on coexistence. Ecological Monographs 92:e01493.

Uriarte, M., and D. Menge. 2018. Variation between individuals fosters regional species coexistence. Ecology Letters 21:1496–1504.

Violle, C., B. J. Enquist, B. J. McGill, L. Jiang, C. H. Albert, C. Hulshof, V. Jung, and J. Messier. 2012. The return of the variance: intraspecific variability in community ecology. Trends in Ecology & Evolution 27:244–252.

Webster, M. M., A. J. W. Ward, and P. J. B. Hart. 2008. Individual boldness affects interspecific interactions in sticklebacks. Behavioral Ecology and Sociobiology 63:511–520.

Wende, B., M. M. Gossner, I. Grass, T. Arnstadt, M. Hofrichter, A. Floren, K. E. Linsenmair, W. W. Weisser, and I. Steffan-Dewenter. 2017. Trophic level, successional age and trait matching determine specialization of deadwood-based interaction networks of saproxylic beetles. Proceedings of the Royal Society B: Biological Sciences 284:20170198.

Wetzel, W. C., H. M. Kharouba, M. Robinson, M. Holyoak, and R. Karban. 2016. Variability in plant nutrients reduces insect herbivore performance. Nature 539:425–427.

Wickman, J., T. Koffel, and C. A. Klausmeier. 2023. A theoretical framework for trait-based eco-evolutionary dynamics: Population structure, intraspecific variation, and community assembly. The American Naturalist 201:501–522.

Witelski, T., and M. Bowen. 2015. Methods of mathematical modelling: continuous systems and differential equations. Springer International Publishing, Cham, CH.

Wood, S. 2006. Generalized additive models: an introduction with R. CRC press, Boca Raton, FL, USA.

