## Supporting information S1-S10 for "Quantifying the effects of intraspecific trait variation and interspecific trait correlations on interacting populations - a nonlinear averaging approach"

This file contains the SI for:

**S4 The Gaussian interaction function** 7

**S5 Approximation when $\gamma$ only depends on the difference in trait value** 9

**S6 Deriving lines of zero and maximal curvature for the Gaussian and the logistic interaction function** 10

S6.1 Gaussian interaction function . . . 10

S6.2 Logistic interaction function . . . 11

**S7 Additional results for the logistic interaction function** 13

**S8 Equilibria and coexistence in the predator-prey model** 13

**S9 Additional results for application example 2** 15

**S10 Application example 3: Lotka-Volterra competition model** 19

### S1 Numerical integration

As an alternative to the sampling method used in the main text, we can also use the pcubature routine from the cubature R package to perform adaptive multidimensional integration of the double integral in Eq. (5) (Narasimhan *et al.*, 2019). This method is used in addition to the sampling method in some of the supplementary figures.

### S2 Derivation of the approximation

#### S2.1 Up to second order

For a two-dimensional function $\gamma(a, b)$, up to second order, the Taylor approximation around the point $(\mu_a, \mu_b)$ is:

$$\begin{aligned}\gamma(a,b) \approx& \gamma(\mu_a,\mu_b) + (a-\mu_a)\left.\frac{\partial\gamma}{\partial a}\right|_{(\mu_a,\mu_b)} + (b-\mu_b)\left.\frac{\partial\gamma}{\partial b}\right|_{(\mu_a,\mu_b)} + \\ &\frac{1}{2}\left((a-\mu_a)^2\left.\frac{\partial^2\gamma}{\partial a^2}\right|_{(\mu_a,\mu_b)} + (b-\mu_b)^2\left.\frac{\partial^2\gamma}{\partial b^2}\right|_{(\mu_a,\mu_b)} + 2(a-\mu_a)(b-\mu_b)\left.\frac{\partial^2\gamma}{\partial a\partial b}\right|_{(\mu_a,\mu_b)}\right).\end{aligned} \tag{S2.1}$$

Taking the expectation over the joint distribution of $a$ and $b$ yields

$$\begin{aligned}\overline{\gamma(a,b)} = \mathbf{E}[\gamma(a,b)] \approx& \mathbf{E}\Bigg[\gamma(\mu_a,\mu_b) + (a-\mu_a)\left.\frac{\partial\gamma}{\partial a}\right|_{(\mu_a,\mu_b)} + (b-\mu_b)\left.\frac{\partial\gamma}{\partial b}\right|_{(\mu_a,\mu_b)} + \\ &\frac{1}{2}\left((a-\mu_a)^2\left.\frac{\partial^2\gamma}{\partial a^2}\right|_{(\mu_a,\mu_b)} + (b-\mu_b)^2\left.\frac{\partial^2\gamma}{\partial b^2}\right|_{(\mu_a,\mu_b)} + 2(a-\mu_a)(b-\mu_b)\left.\frac{\partial^2\gamma}{\partial a\partial b}\right|_{(\mu_a,\mu_b)}\right)\Bigg].\end{aligned} \tag{S2.2}$$

We can take the expectation for each term independently. Furthermore, the partial derivatives can be taken in front of the sum, since these are always approximated at the same point and do not depend on $a$ and $b$. Moreover

$$\mathbf{E}[(a-\mu_a)] = 0 \tag{S2.3}$$

$$\mathbf{E}[(a-\mu_a)^2] = \sigma_a^2 \tag{S2.4}$$

$$\mathbf{E}[(a-\mu_a)(b-\mu_b)] = \sigma_{a,b}, \tag{S2.5}$$

where $\sigma_{a,b}$ is the covariance between $a$ and $b$. Finally, we find that:

$$\overline{\gamma(a,b)} \approx \gamma(\mu_a,\mu_b) + \frac{1}{2}\sigma_a^2\left.\frac{\partial^2\gamma}{\partial a^2}\right|_{(\mu_a,\mu_b)} + \frac{1}{2}\sigma_b^2\left.\frac{\partial^2\gamma}{\partial b^2}\right|_{(\mu_a,\mu_b)} + \sigma_{a,b}\left.\frac{\partial^2\gamma}{\partial a\partial b}\right|_{(\mu_a,\mu_b)}. \tag{S2.6}$$

Where we note that the trait covariance can be expressed as the product of an independent trait correlation coeffficent ($\rho_{a,b}$) that can range from -1 to 1 and the single species trait standard deviations: $\sigma_{a,b} = \rho_{a,b}\sigma_a\sigma_b$.

#### S2.2 Third order approximation for the multivariate normal distribution

In this section we will extend the approximation to higher order for cases where $a$ and $b$ are generated by a bivariate normal distribution. According to Isserlis' theorem (Isserlis, 1918), in this case the third order central moments should all be zero and hence third-order derivatives do not contribute to an approximation of $\overline{\gamma(a,b)}$. For traits that are distributed according to a bivariate normal distribution our second order approximation is therefore actually valid up to third order.

#### S2.3 Higher order approximations

Using Isserlis' theorem, it is possible to calculate higher-order approximations as well for traits that are distributed according to a bivariate normal distribution. The approximation of the order of $\kappa$ can be written as:

$$\gamma(a,b) \approx \gamma(\mu_a,\mu_b) + \sum_{d=1}^{\kappa} \left( \frac{1}{d!} \sum_{n_\mathrm{a}=0}^{d} \binom{d}{n_\mathrm{a}} (a-\mu_a)^{n_\mathrm{a}} (b-\mu_b)^{d-n_\mathrm{a}} \left. \frac{\partial^d \gamma}{\partial a^{n_\mathrm{a}} \partial b^{d-n_\mathrm{a}}} \right|_{\mu_a,\mu_b} \right). \tag{S2.7}$$

Thus

$$\overline{\gamma(a,b)} \approx \gamma(\mu_a,\mu_b) + \sum_{d=1}^{\kappa} \left( \frac{1}{d!} \sum_{n_\mathrm{a}=0}^{d} \binom{d}{n_\mathrm{a}} \mathbf{E}[(a-\mu_a)^{n_\mathrm{a}} (b-\mu_b)^{d-n_\mathrm{a}}] \left. \frac{\partial^d \gamma}{\partial a^{n_\mathrm{a}} \partial b^{d-n_\mathrm{a}}} \right|_{\mu_a,\mu_b} \right). \tag{S2.8}$$

Isserlis' theorem allows us to calculate $\mathbf{E}[(a-\mu_a)^{n_\mathrm{a}}(b-\mu_b)^{d-n_\mathrm{a}}]$ for bivariate normal distributions. Specifically, it states that $\mathbf{E}[(a-\mu_a)^{n_\mathrm{a}}(b-\mu_b)^{d-n_\mathrm{a}}] = 0$ whenever $d$ is odd. For the even cases, it states:

$$\mathbf{E}[(a-\mu_a)^{n_\mathrm{a}}(b-\mu_b)^{d-n_\mathrm{a}}] = \sum \sigma_a^{2^x} \sigma_b^{2^y} \sigma_{a,b}^z. \tag{S2.9}$$

The sum is taken over all possible ways of partitioning a set of $n_\mathrm{a}$ $a-\mu_a$ items and $d-n_\mathrm{a}$ $b-\mu_b$ items into $\frac{d}{2}$ pairs. For each partition, $x$ corresponds to the number $a-\mu_a$,$a-\mu_a$ pairs it contains, $y$ to the number of $b-\mu_b$,$b-\mu_b$ pairs it contains and $z$ to the number of $a-\mu_a$,$b-\mu_b$ pairs with $x+y+z=\frac{d}{2}$. Given $n_\mathrm{a}$, the number of pairs of each type is fully specified by the number of mixed pairs, i.e. $x=\frac{1}{2}(n_\mathrm{a}-z)$ and $y=\frac{d-n_\mathrm{a}-z}{2}$. Thus the sum can be formulated to run over 1 dimension only:

$$\mathbf{E}[(a-\mu_a)^{n_\mathrm{a}}(b-\mu_b)^{d-n_\mathrm{a}}] = \sum_{z=z_0}^{\min(n_\mathrm{a},d-n_\mathrm{a})} N_z \sigma_a^{2^{\frac{1}{2}(n_\mathrm{a}-z)}} \sigma_b^{2^{\frac{1}{2}(d-n_\mathrm{a}-z)}} \sigma_{a,b}^z, \tag{S2.10}$$

with

$$z_0 = \begin{cases} 0 & \text{if } n_\mathrm{a} \text{ is even} \\ 1 & \text{if } n_\mathrm{a} \text{ is odd} \end{cases} \tag{S2.11}$$

and $N_z$ the number of partitions with $z$ mixed $a-\mu_a$, $b-\mu_b$ pairs.

The accuracy of the higher order approximations for a Gaussian interaction function depends on the amount of intraspecific variation (figure S2.1). This figure shows that the sixth and fourth order approximations are closer to the exact sampled values than the standard second order approximation when $\sigma_a$ is small. However, as the amount of variation increases, the higher order approximations diverge more strongly from the exact values, until they become even less accurate than the naive values. Additionally, note that the the direction of the divergence is inverted from one approximation to the next higher even order.

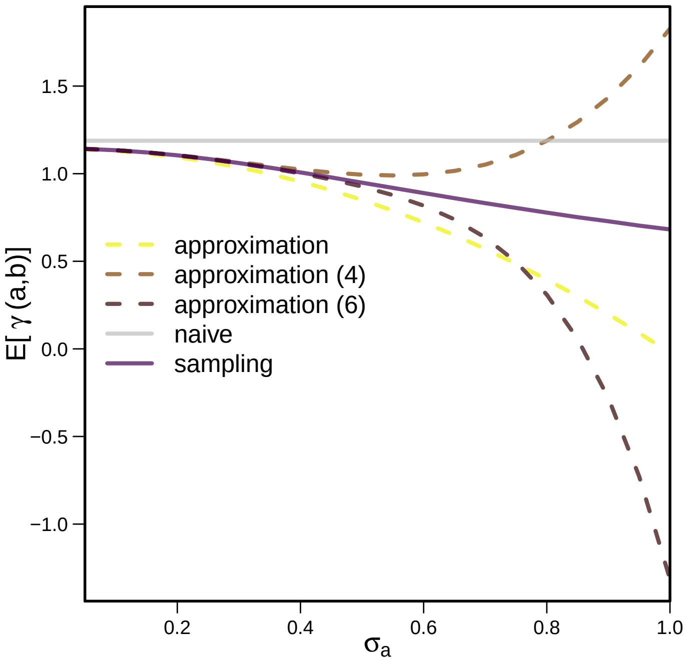

**Figure S2.1** – Estimates of $\gamma$ using the Gaussian interaction function with $c_{max} = 1.2$, $\omega = 1$, $\mu_a = 0$, $\mu_b = 0.1$, $\rho_{a,b} = 0$, $\sigma_b = 0.2$.

### S3 The polynomial interaction function

#### S3.1 Covariance and the direction of the effect of ITV on the average interaction value

Here we consider the case where $p_1$ and $p_2$ in the polynomial interaction function have the same sign. If both are positive and $\sigma_{a,b} = 0$, intraspecific variation in either species always has a positive effect. The analogous result is true if both $p_1$ and $p_2$ are negative. Here we explore under what conditions the direction of the effect holds even with a non-zero covariance and under what conditions a non-zero covariance can change the direction of the effect.

Assume that $p_1, p_2 > 0$. Then, according to Eq. (8), to find conditions for $p_3$ such that the effect of ITV will be positive irrespective of the parameters of the joint distribution, we are looking for the values of $p_3$ such that

$$p_1\sigma_a^2 + p_2\sigma_b^2 + p_3\sigma_{a,b} > 0 \tag{S3.1}$$

for all possible bivariate distributions. We rewrite this as:

$$(\sqrt{p_1}\sigma_a - \sqrt{p_2}\sigma_b)^2 + 2\sqrt{p_1}\sqrt{p_2}\sigma_a\sigma_b + p_3\sigma_{a,b} > 0. \tag{S3.2}$$

Since the minimum value of the first term across all possible joint distributions is zero (we can choose $\sigma_b = (\sqrt{p_1}/\sqrt{p_2})\sigma_a$), for the effect to be positive for all choices of $\sigma_a$, $\sigma_b$, and $\sigma_{a,b}$, we need the sum of the remaining terms to be positive:

$$p_3\sigma_{a,b} + 2\sqrt{p_1}\sqrt{p_2}\sigma_a\sigma_b > 0 \tag{S3.3}$$

$$-p_3\frac{\sigma_{a,b}}{\sigma_a\sigma_b} < 2\sqrt{p_1}\sqrt{p_2}. \tag{S3.4}$$

By the Cauchy-Schwarz inequality, $-\sigma_a\sigma_b \leq \sigma_{a,b} \leq \sigma_a\sigma_b$ and thus the term on the left ranges between $p_3$ and $-p_3$. The condition must hold over this whole range, and therefore we must have $|p_3| < 2\sqrt{p_1}\sqrt{p_2}$ so that the effect of intraspecific trait variation is always positive. Analogously if $p_1, p_2 < 0$, the effect of intraspecific trait variation is always negative if $|p_3| < 2\sqrt{-p_1}\sqrt{-p_2}$.

#### S3.2 Imposing limits on the polynomial interaction function

We now apply our approach to a complex, *non-differentiable* interaction function, that is inspired by a system where predators feed on prey that may be toxic. For example, the ability of snakes to feed on cane toads depends on the body size of the snake, which determines its susceptibility to toxin, and the body size of the toad, which predicts its toxicity (Phillips *et al.*, 2003; Phillips & Shine, 2004). We assume selection has acted on the gape width, such that snakes cannot ingest a prey item that would be lethal to them. As a consequence, the rate of successful attack $\gamma$ depends on the toxicity of the prey ($a$) and the resistance to toxicity of the predator ($b$):

$$\gamma(a,b) = \left((p_{\max} - c_1 b^+) - (d_{\max} - c_2 b^+)^+ {a^+}^2\right)^+, \tag{S3.5}$$

with $x^+ \equiv \max(x, 0)$. The actual predation rate is $p_{\max}$ at most, and decreases with the squared toxicity of the prey ($a^2$) multiplied with a sensitivity factor. This sensitivity factor has a maximum of $d_{\max}$, but decreases with predator toxin resistance, by an amount $c_2 b$. However, the maintenance of the resistance leads to a lower overall predation efficiency (by an amount $c_1 b$). As a consequence, defense against toxicity only pays off when toxic prey is encountered. Furthermore, any negative values of $a$, $b$, $\gamma(a,b)$ and $d_{\max} - c_2 b$ are treated as zero, to ensure biologically realistic behavior. This leads to an interaction function that is non-differentiable, for example when $d_{\max} = c_2 b$ or $b = 0$. Around these lines, the derivatives and curvatures are discontinuous.

**Table S3.1** – Overview of the non-differential interaction function and its second order partial derivatives that were used in Eq. 6 to approximate $\overline{\gamma(a,b)}$. For this interaction function $\gamma(a,b)$, $a$, $b$, and $d_{\max} - c_2 b$ are forced to be non-negative. The effect of this procedure depends on the order of checking the terms and setting them to zero, in our case we first test $a$, $b$ and $d_{\max} - c_2 b$, and $\gamma(a,b)$ itself is checked last. In the table, two cases are shown: first the case is shown where none of the checked values was below zero, and on the second row the case where $d_{\max} - c_2 b$ was initially below zero.

| $\gamma(a,b)$ | $\frac{\partial^2\gamma}{\partial a^2}$ | $\frac{\partial^2\gamma}{\partial b^2}$ | $\frac{\partial^2\gamma}{\partial a\partial b}$ |
|---|---|---|---|
| $p_{\max} - c_1 b - (d_{\max} - c_2 b)a^2$ | $-2(d_{\max} - c_2 b)$ | 0 | $2c_2 a$ |
| $p_{\max} - c_1 b$ | 0 | 0 | 0 |

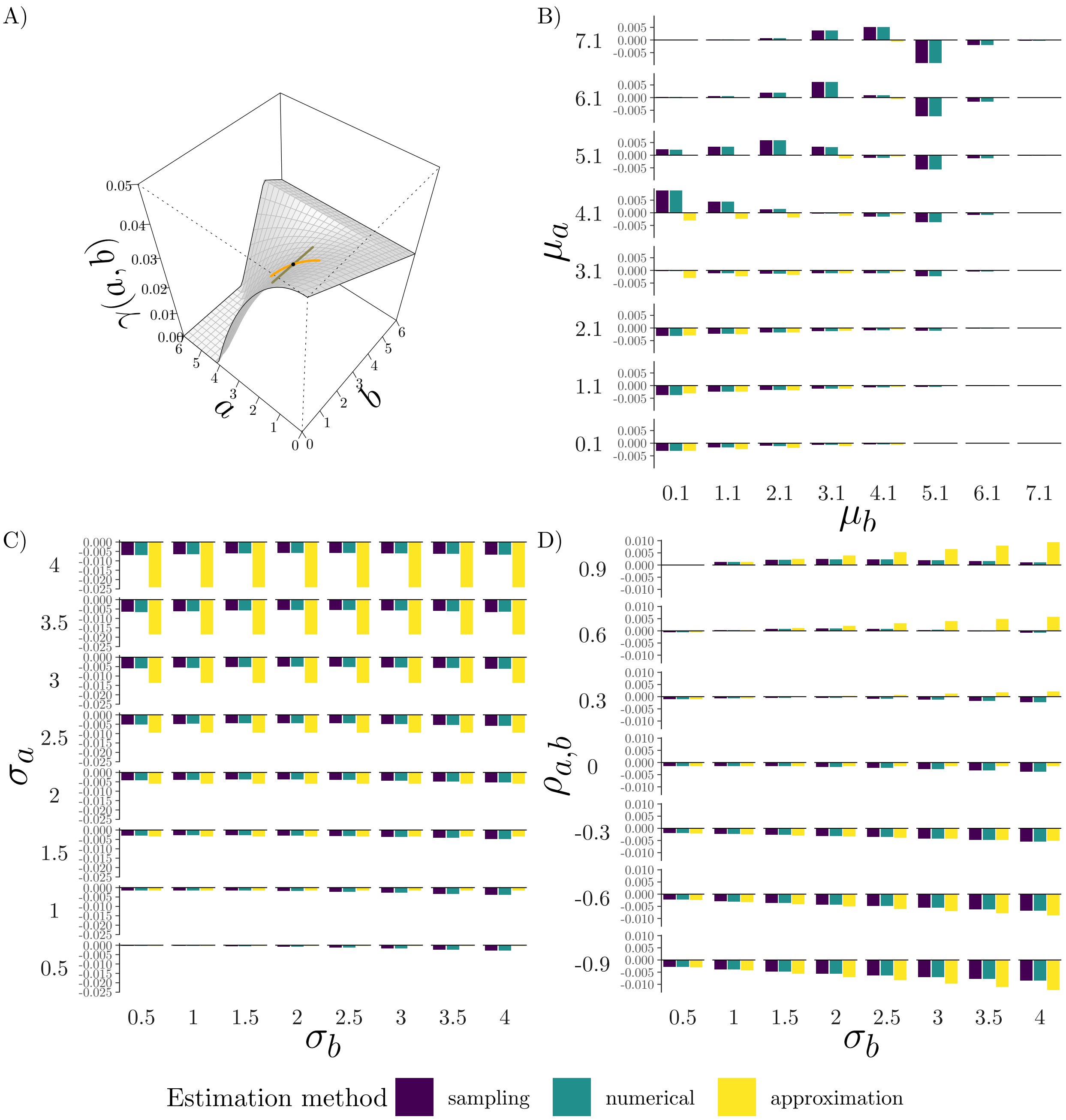

**Figure S3.1** – Effect of ITV on the non-differentiable interaction function. A) illustration of the interaction function. B-D) Difference between the naive estimate of the average of the non-differentiable interaction function and the value of three estimates that take ITV into account. $p_{\max} = 0.05$, $d_{\max} = 0.003$, $c_1 = 0.004$ and $c_2 = 0.0006$, at $\mu_a = 2.5$, $\mu_b = 2.5$. Unless stated differently on the axes, $\mu_a$ and $\mu_b$ are at their respective reference values (represented by the cross), $\sigma_a = \sigma_b = 1$ and $\sigma_{a,b} = 0$.

In our example, the exact estimates show strong effects of ITV when trait means are close to the discontinuities (figure S3.1B). However, in such cases the approximation performs poorly, because it cannot capture dynamics across a discontinuity. Further away from the discontinuities, the approximation again performs satisfactorily. In particular, when predators are very large, to the extent that they are completely immune to the toxin, the interaction function is linear due to the cost of resistance. As a consequence, trait variation does not affect the average interaction here, and hence the naive value and the other estimates are identical (figure S3.1B, $\mu_b = 7.1$). Because the system contains only linear terms in $b$, variation in $b$ has direct effects only through the discontinuities (which cannot be captured by the approximation, figure S3.1C). However, variation in $b$ can have indirect effects, also on the approximation, through covariance with the prey trait, which can sometimes be strong enough to overpower the direct effect of variation in $a$ (figure S3.1D). Non-differentiable interaction functions such as this one may occur in nature when there are hard bounds on the interaction function. Even if such limits are approached more smoothly, large changes in curvature may occur within a very small trait value range and the approximation of the effect of ITV in such a region would be inaccurate, as for a non-differentiable function.

### S4 The Gaussian interaction function

As an additional example to the polynomial and logistic interaction functions considered in section 2.3, we explore the *Gaussian interaction function*. This function is commonly used in ecological and evolutionary models, for example to model the attack rate of predators or parasites, the strength of competition, or the strength of mutualisms that depend on matching traits (e.g. Gibert & Brassil, 2014; Barabás & D'Andrea, 2016; Hart *et al.*, 2016; Senthilnathan & Gavrilets, 2021). Like the logistic interaction function, it can be written in terms of the differences in trait values. The strength of the interaction decreases with increasing trait difference between the interacting individuals:

$$\gamma(a,b) = c_{\max} e^{\left(-\frac{(a-b)^2}{\omega^2}\right)}, \tag{S4.1}$$

where $c_{\max}$ is the maximum possible interaction coefficient and $\omega$ the interaction width of the Gaussian curve (figure S4.1A). The second derivatives with respect to the two traits are equal

$$\frac{\partial^2\gamma}{\partial a^2} = \frac{\partial^2\gamma}{\partial b^2} = \frac{4(a-b)^2 - 2\omega^2}{\omega^4}\gamma(a,b). \tag{S4.2}$$

Thus, according to the analytic approximation Eq. (6) and as expected for a function that only depends on the difference in trait value, $\sigma_a$ and $\sigma_b$ have identical effects on $\overline{\gamma(a,b)}$. Furthermore,

$$\frac{\partial^2\gamma}{\partial a \partial b} = -\frac{\partial^2\gamma}{\partial a^2}. \tag{S4.3}$$

When the difference in trait values is small compared to the interaction width $\omega$, the curvature is negative (around the blue line in figure S4.1A). Thus, when the species' trait means are close together, the $\overline{\gamma(a,b)}$ values that take into account intraspecific variation are lower than the naive estimate (dark orange and pink bars in figure S4.1B). Intuitively this occurs because variation introduces individuals that are more different from the bulk of individuals in both species $A$ and $B$, thereby reducing the average similarity between the species. However, when trait values lie further apart, curvatures are positive (around the red line in figure S4.1A), and the effect of ITV is reversed (right three bars in figure S4.1B). Here, ITV increases similarity and thereby average interaction strength. In both cases, the impact of ITV becomes stronger when the traits of the two species are negatively correlated and weaker if they are positively correlated. With increasing $\omega$, the interaction becomes less trait-dependent and values of $\overline{\gamma(a,b)}$ approach $\gamma(\mu_a, \mu_b)$, as is evident from the vanishing curvatures as $\omega$ goes to infinity (Eq. (S4.2)). At this limit, interaction strength is maximal, since $\lim_{\omega\to\infty} \gamma(a,b) = c_{\max}$.

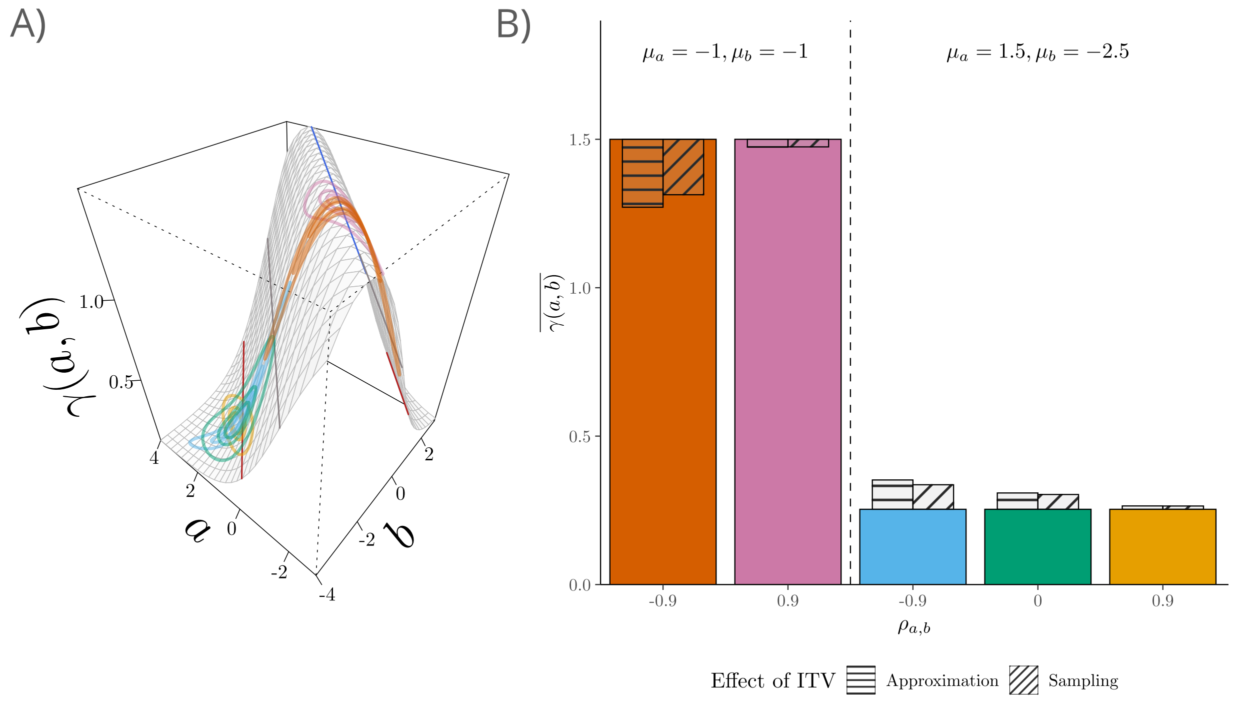

**Figure S4.1** – Effect of ITV on the average interaction parameter ($\overline{\gamma(a,b)}$) for the Gaussian interaction function Eq. (S4.1). A: Illustration of the Gaussian interaction function $\gamma$ with $c_{\max} = 1.5$ and $\omega = 3$ with lines of zero (grey), maximal (dark red) and minimal curvature (blue). B: The colored bars give the average interaction parameter without ITV or equivalently the naive estimate of the average interaction parameter $\gamma(\mu_a, \mu_b)$. The striped and hatched bars represent how ITV changes the average interaction parameter compared to the naive case/the case without ITV. The joint trait distributions used in B are shown as ellipses A. For each distribution three ellipses are shown that contain, from inside to outside, 25%, 50% and 90% of the interactions for that distribution. $\sigma_a = 0.75$, $\sigma_b = 0.5$.

### S5 Approximation when $\gamma$ only depends on the difference in trait value

A special case emerges when $\gamma(a, b)$ only depends on the difference between $a$ and $b$ and hence can be written as $\gamma(u(a, b))$, with $u(a, b) = a - b$. Using the chain rule, we can find the first order partial derivatives for $\gamma$:

$$\frac{\partial \gamma}{\partial a} = \frac{\partial \gamma}{\partial u}\frac{\partial u}{\partial a} \tag{S5.1}$$

$$\frac{\partial \gamma}{\partial b} = \frac{\partial \gamma}{\partial u}\frac{\partial u}{\partial b}, \tag{S5.2}$$

given the definition of $u$, this becomes:

$$\frac{\partial \gamma}{\partial a} = \frac{\partial \gamma}{\partial u} \tag{S5.3}$$

$$\frac{\partial \gamma}{\partial b} = -\frac{\partial \gamma}{\partial u}. \tag{S5.4}$$

Hence, in this case we find that $\frac{\partial \gamma}{\partial a} = -\frac{\partial \gamma}{\partial b}$. We proceed to calculate the second order derivatives, where we use the symmetry of derivatives ($\frac{\partial}{\partial x}\left(\frac{\partial f}{\partial y}\right) = \frac{\partial}{\partial y}\left(\frac{\partial f}{\partial x}\right)$) and Eq. S5.3–S5.4:

$$\frac{\partial^2 \gamma}{\partial a^2} = \frac{\partial}{\partial a}\left(\frac{\partial \gamma}{\partial u}\right) = \frac{\partial}{\partial u}\left(\frac{\partial \gamma}{\partial a}\right) = \frac{\partial}{\partial u}\left(\frac{\partial \gamma}{\partial u}\right) \tag{S5.5}$$

$$\frac{\partial^2 \gamma}{\partial b^2} = -\frac{\partial}{\partial b}\left(\frac{\partial \gamma}{\partial u}\right) = -\frac{\partial}{\partial u}\left(\frac{\partial \gamma}{\partial b}\right) = \frac{\partial}{\partial u}\left(\frac{\partial \gamma}{\partial u}\right) \tag{S5.6}$$

$$\frac{\partial^2 \gamma}{\partial a \partial b} = \frac{\partial}{\partial b}\left(\frac{\partial \gamma}{\partial u}\right) = \frac{\partial}{\partial u}\left(\frac{\partial \gamma}{\partial b}\right) = -\frac{\partial}{\partial u}\left(\frac{\partial \gamma}{\partial u}\right). \tag{S5.7}$$

This result shows that when $\gamma(a, b)$ only depends on the difference between $a$ and $b$,

$$\frac{\partial^2 \gamma}{\partial a^2} = \frac{\partial^2 \gamma}{\partial b^2} = -\frac{\partial^2 \gamma}{\partial a \partial b}. \tag{S5.8}$$

We can use this identity to simplify the approximation of $\overline{\gamma(a, b)}$ from Eq. 6:

$$\overline{\gamma(a, b)} \approx \gamma(\mu_a, \mu_b) + \left.\frac{\partial^2 \gamma}{\partial a^2}\right|_{(\mu_a, \mu_b)} \left(\frac{1}{2}\sigma_a^2 + \frac{1}{2}\sigma_b^2 - \sigma_{a,b}\right). \tag{S5.9}$$

Hence, an increase in variance in $a$ has the exact same effect as an increase in variance in $b$. Furthermore, increasing covariance between $a$ and $b$ acts opposed to the direct effect of the variances. However, by the Cauchy-Schwarz inequality, $|\sigma_{a,b}| \leq \sigma_a \sigma_b$ and thus

$$\left(\frac{1}{2}\sigma_a^2 + \frac{1}{2}\sigma_b^2 - \sigma_{a,b}\right) \geq \frac{1}{2}\left(\sigma_a^2 - 2\sigma_a\sigma_b + \sigma_b^2\right) = \frac{1}{2}\left(\sigma_a - \sigma_b\right)^2 \geq 0. \tag{S5.10}$$

Hence the covariance term can never change the sign of the overall effect. In the special case when $a$ and $b$ have the same variance and they are perfectly correlated ($\rho_{a,b} = 1 \rightarrow \sigma_{a,b} = \sigma_a\sigma_b$), the term on the right cancels out and the expression can be simplified to:

$$\overline{\gamma(a, b)} \approx \gamma(\mu_a, \mu_b). \tag{S5.11}$$

### S6 Deriving lines of zero and maximal curvature for the Gaussian and the logistic interaction function

#### S6.1 Gaussian interaction function

All curvatures of the Gaussian interaction function (see Eq. (S4.2)) vanish when:

$$\frac{4(a-b)^2 - 2\omega^2}{\omega^4}\gamma(a,b) = 0. \tag{S6.1}$$

This occurs when either $\gamma(a,b) = 0$, and hence when the difference between $a$ and $b$ goes to infinity, or when:

$$4(a-b)^2 - 2\omega^2 = 0, \tag{S6.2}$$

or:

$$a - b = \pm\frac{\omega}{\sqrt{2}}. \tag{S6.3}$$

The effect of variation is the strongest (at least for small variance and in the approximation) when the curvature reaches an extreme value. In the $a$ direction, this occurs when:

$$\frac{\partial^3\gamma(a,b)}{\partial a^3} = 0, \tag{S6.4}$$

or:

$$\frac{\partial}{\partial a}\left(\frac{4(a-b)^2 - 2\omega^2}{\omega^4}\gamma(a,b)\right) = 0, \tag{S6.5}$$

which is equal to:

$$\left(\frac{4(a-b)^2 - 2\omega^2}{\omega^4}\right)\frac{\partial\gamma(a,b)}{\partial a} + \gamma(a,b)\frac{\partial}{\partial a}\left(\frac{4(a-b)^2 - 2\omega^2}{\omega^4}\right) = 0. \tag{S6.6}$$

This expression can be simplified to:

$$\gamma(a,b)(a-b)\left(\frac{4(a-b)^2 - 2\omega^2}{\omega^4}(\frac{-2}{\omega^2}) + \frac{8}{\omega^4}\right) = 0. \tag{S6.7}$$

This occurs when either $\gamma(a,b) = 0$ (when the absolute difference between $a$ and $b$ goes to infinity), when $a = b$ and when:

$$\frac{4(a-b)^2}{\omega^4} = \frac{6}{\omega^2}. \tag{S6.8}$$

or:

$$(a-b)^2 = 1.5\omega^2, \tag{S6.9}$$

or:

$$a - b = \pm\sqrt{3}\frac{\omega}{\sqrt{2}}. \tag{S6.10}$$

The earlier found line at $a = b$ corresponds to a line of minimal (negative) curvature, while this second line (Eq. S6.10) corresponds to a line of maximal curvature.

##### S6.2 Logistic interaction function

For the logistic interaction function

$$\gamma(a,b) = p_{\max}\frac{1}{1+e^{-k(b-a-h)}}, \tag{S6.11}$$

all curvatures (see Table 1) vanish when:

$$k^2\left(\gamma(a,b) + \frac{2\gamma(a,b)^3}{p_{\max}^2} - 3\frac{\gamma(a,b)^2}{p_{\max}}\right) = 0. \tag{S6.12}$$

With $k \neq 0$, this occurs when:

$$\gamma(a,b)\left(1 + \frac{2\gamma(a,b)^2}{p_{\max}^2} - 3\frac{\gamma(a,b)}{p_{\max}}\right) = 0. \tag{S6.13}$$

This occurs either when $b-a$ goes to minus infinity and hence $\gamma(a,b) = 0$, or when:

$$1 + \frac{2\gamma(a,b)^2}{p_{\max}^2} - 3\frac{\gamma(a,b)}{p_{\max}} = 0. \tag{S6.14}$$

Which can be solved straightforwardly using the quadratic formula:

$$\gamma(a,b) = \frac{\frac{3}{p_{\max}} \pm \sqrt{\frac{9}{p_{\max}^2} - \frac{8}{p_{\max}^2}}}{\frac{4}{p_{\max}^2}} \tag{S6.15}$$

or simplified:

$$\gamma(a,b) = \frac{(3\pm 1)p_{\max}}{4}. \tag{S6.16}$$

We use the definition of $\gamma$ to solve for a relation between $a$ and $b$:

$$p_{\max}\frac{1}{1+e^{-k(b-a-h)}} = \frac{(3\pm 1)p_{\max}}{4}, \tag{S6.17}$$

or:

$$1+e^{-k(b-a-h)} = \frac{4}{3\pm 1}, \tag{S6.18}$$

Because $\frac{4}{3\pm 1} = 1.5 \pm 0.5$, this is equivalent to

$$e^{-k(b-a-h)} = 0.5 \pm 0.5. \tag{S6.19}$$

This occurs when either $b-a$ goes to infinity, and so the left side goes to zero, or when the left side is equal to one, which corresponds to:

$$-k(b-a-h) = 0. \tag{S6.20}$$

Hence the line at which the curvature vanishes lies at:

$$b - a = h. \tag{S6.21}$$

In contrast, the values for $\overline{a}$ and $\overline{b}$ where the approximation predicts the largest difference to the naive estimate, occurs when the curvature reaches an extreme value. We start by calculating the maximal curvature in the $a$ direction. We do so by setting the third order derivative to zero:

$$\frac{\partial^3\gamma(a,b)}{\partial a^3} = \frac{\partial}{\partial a}\frac{\partial^2\gamma(a,b)}{\partial a^2} = \frac{\partial}{\partial a}k^2(\gamma(a,b) + \frac{2\gamma(a,b)^3}{p_{\max}^2} - 3\frac{\gamma(a,b)^2}{p_{\max}}) = 0, \tag{S6.22}$$

which is equivalent to:

$$\frac{\partial\gamma(a,b)}{\partial a} + \frac{6\gamma(a,b)^2}{p_{\max}^2}\frac{\partial\gamma(a,b)}{\partial a} - 6\frac{\gamma(a,b)}{p_{\max}}\frac{\partial\gamma(a,b)}{\partial a} = 0. \tag{S6.23}$$

As a consequence, either

$$\frac{\partial\gamma(a,b)}{\partial a} = 0, \tag{S6.24}$$

which occurs when $\overline{a} \to \infty$ or $\overline{b} \to \infty$, or:

$$1 + \frac{6}{(1+e^{-k(b-a-h)})^2} - \frac{6}{1+e^{-k(b-a-h)}} = 0. \tag{S6.25}$$

We simplify this equation by defining $\eta = 1 + e^{-k(b-a-h)}$:

$$1 + \frac{6}{\eta^2} - \frac{6}{\eta} = 0. \tag{S6.26}$$

This can be written as a standard quadratic equation:

$$\eta^2 + 6 - 6\eta = 0. \tag{S6.27}$$

This is equivalent to:

$$(\eta - 3)^2 - 3 = 0, \tag{S6.28}$$

and hence:

$$\eta = 3 \pm \sqrt{3}. \tag{S6.29}$$

Putting the definition of $\eta$ back in and solving for $a$ gives:

$$a = b - h + \frac{\ln(2 \pm \sqrt{3})}{k}. \tag{S6.30}$$

To show that the extrema are symmetric around the inflection line $a = b - h$ (see Eq. S6.21), we note that:

$$2 - \sqrt{3} = \frac{2-\sqrt{3}}{4-3} = \frac{2-\sqrt{3}}{(2+\sqrt{3})(2-\sqrt{3})} = \frac{1}{2+\sqrt{3}} = (2+\sqrt{3})^{-1}. \tag{S6.31}$$

Since $\ln(x^{-1}) = -\ln(x)$ we can therefore write:

$$a = b - h \pm \frac{\ln(2 + \sqrt{3})}{k}. \tag{S6.32}$$

or:

$$b = a + h \pm \frac{\ln(2 + \sqrt{3})}{k}. \tag{S6.33}$$

From Eq. 10, we can deduce the difference between the approximated value of $\gamma$ and the naive value as:

$$\overline{\gamma(a,b)} - \gamma(\mu_a,\mu_b) = \left.\frac{\partial^2\gamma}{\partial a^2}\right|_{(\mu_a,\mu_b)}\left(\frac{1}{2}\sigma_a^2 + \frac{1}{2}\sigma_b^2 - \sigma_{a,b}\right) \tag{S6.34}$$

From this expression it is clear to see that when the trait variances and covariance are kept constant, the largest difference occurs when the curvature reaches an extreme value, which in turn occurs when:

$$\mu_a = \mu_b - h \pm \frac{\ln(2+\sqrt{3})}{k}, \tag{S6.35}$$

as derived above. Whether the line corresponds to maximum or minimum curvature depends on the sign of the $\pm$.

### S7 Additional results for the logistic interaction function

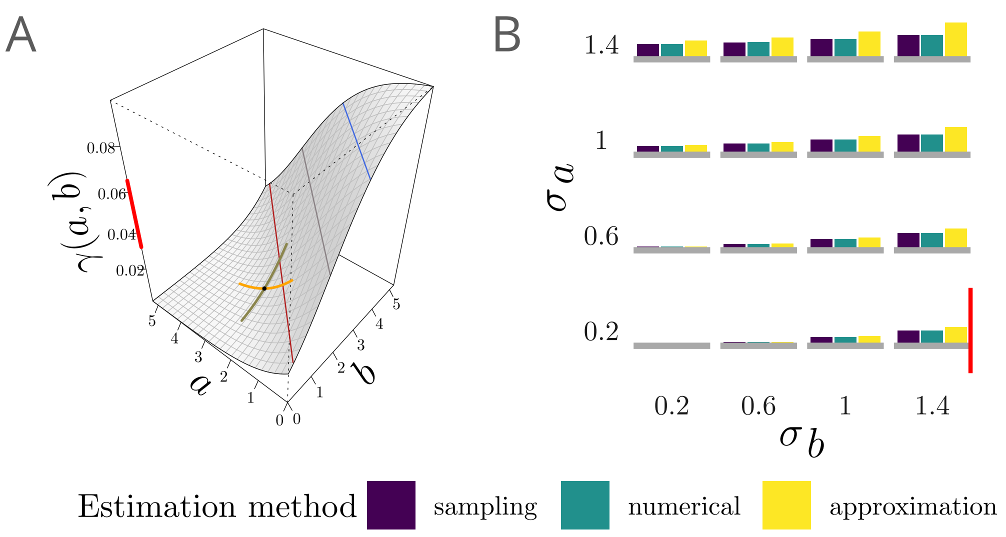

**Figure S7.1** – Effect of increasing trait variances on the effect of ITV and the accuracy of the approximation for the logistic interaction function with $p_{\max} = 0.1$, $k = 1$, $h = 1.5$ at $\mu_a = 2.5$, $\mu_b = 2$ (A). Lines of zero (grey), maximal (dark red) and minimal curvature (blue) are shown. (B) Difference between the various estimates of $\overline{\gamma(a,b)}$ and the naive estimate $\gamma(\mu_a, \mu_b)$ ($\overline{\gamma(a,b)} - \gamma(\mu_a,\mu_b)$) for joint trait distributions with different combinations of trait standard deviations while the mean is kept at the black point in A. The scale of the bars relative to the scale of the interaction function plot in the left column is indicated by bright red vertical line of constant length across panels A and B.

### S8 Equilibria and coexistence in the predator-prey model

The system:

$$\begin{aligned}\frac{\mathrm{d}N_A}{\mathrm{d}t} &= r_A N_A \left(1 - \frac{N_A}{K}\right) - N_A N_B \phi \\ \frac{\mathrm{d}N_B}{\mathrm{d}t} &= r_B N_B + c N_A N_B \phi,\end{aligned} \tag{S8.1}$$

has three equilibrium points $(N_A^* = 0, N_B^* = 0), (N_A^* = K, N_B^* = 0)$ and $(N_A^* = \frac{-r_B}{c\phi}, N_B^* = \frac{r_A}{\phi}(1 + \frac{r_B}{c\phi K}))$, where the third is the potential coexistence equilibrium where both species have non-zero abundance.

To determine whether coexistence is possible, we determine when prey have a positive per-capita growth rate when starting from zero abundance, with predators at their equilibrium value in the absence of prey (i.e.

$N_B^* = 0$):

$$r_A > 0, \tag{S8.2}$$

and similarly when the predators have a positive per-capita growth rate when at zero abundance, with prey at their equilibrium value in the absense of predators (i.e. $N_A^* = K$):

$$r_B + cK\phi > 0. \tag{S8.3}$$

Since $r_A$ is always positive (convention), the condition for coexistence becomes:

$$\phi > -\frac{r_B}{cK}. \tag{S8.4}$$

Changes in $\phi$ due to ITV may thus affect the coexistence status of the two species in either direction, depending on the threshold location and the direction and magnitude of the effect of ITV on $\phi$. For example a decrease in $\phi$ below the threshold in value in Eq. S8.4 leads to predator extinction making the second equilibrium ($N_A^* = K, N_B^* = 0$) asymptotically stable as the remaining equilibria becomes unstable.

The equilibrium abundances change with $\phi$ as evidenced by their derivatives:

$$\frac{\mathrm{d}N_A^*}{\mathrm{d}\phi} = \frac{r_B}{c\phi^2} < 0 \quad \text{for } r_B < 0 \tag{S8.5}$$

$$\frac{\mathrm{d}N_B^*}{\mathrm{d}\phi} = -\frac{r_A}{\phi^2}(1 + \frac{2r_B}{c\phi K}). \tag{S8.6}$$

By setting Eq. S8.6 equal to zero, it can be shown that the equilibrium predator abundance has an extremum at $\hat{\phi} = -\frac{2r_B}{cK}$. Here, $N_B^* = -\frac{r_A cK}{4r_B}$. The value of the second derivative at $\hat{\phi}$ ($-r_A\hat{\phi}^{-3}$) is negative and thus the equilibrium abundance is a maximum. The corresponding value of $N_A^* = \frac{K}{2}$, is half the carrying capacity for the prey species.

### S9 Additional results for application example 2

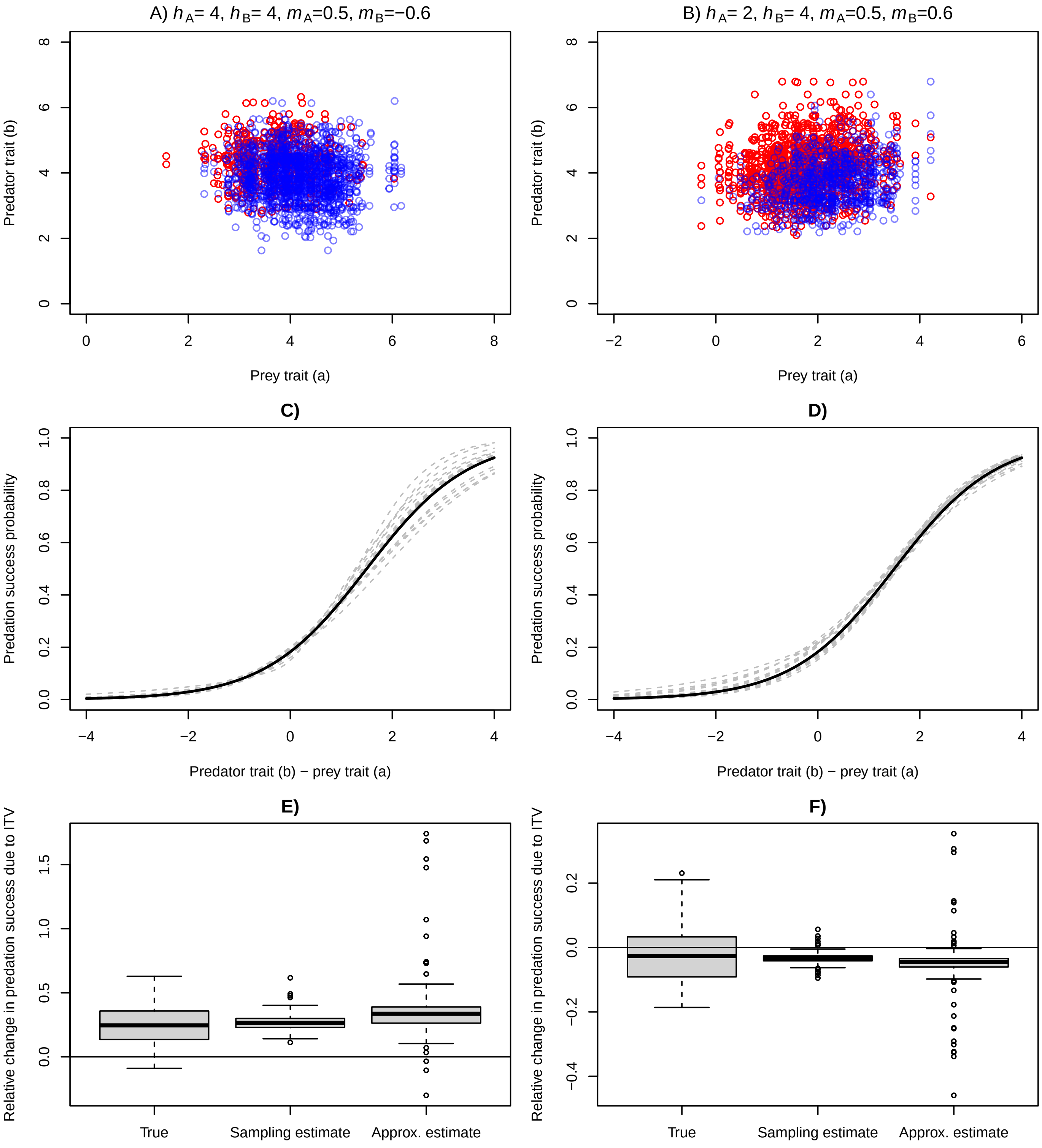

**Figure S9.1** – Estimating the impact of ITV on average predation success from field data with a smaller environmental variance $\sigma_e = 0.5$ compared to $\sigma_e = 1.5$ in Fig. 5. The two columns show two scenarios that differ in the reaction norms via which individuals respond to their micro-environment. A, B) Trait values of pairs of interacting individuals in the first replicate, with successful kills shown in red and encounters where the prey escaped shown in blue. Across all replicates in A, there were on average 2191.5 encounters, with 500.7 kills and 1690.8 escapes. In B, there were on average 2187.7 encounters, with 1318.2 kills and 869.5 escapes. C, D) Estimated interaction function for the first 20 replicates shown in dashed gray lines. Note that we have evaluated the two-dimensional interaction function at the respective mean prey trait $h_A$. The true interaction function is shown in black. E, F) Boxplots of true and estimated effects of ITV on total predation success.

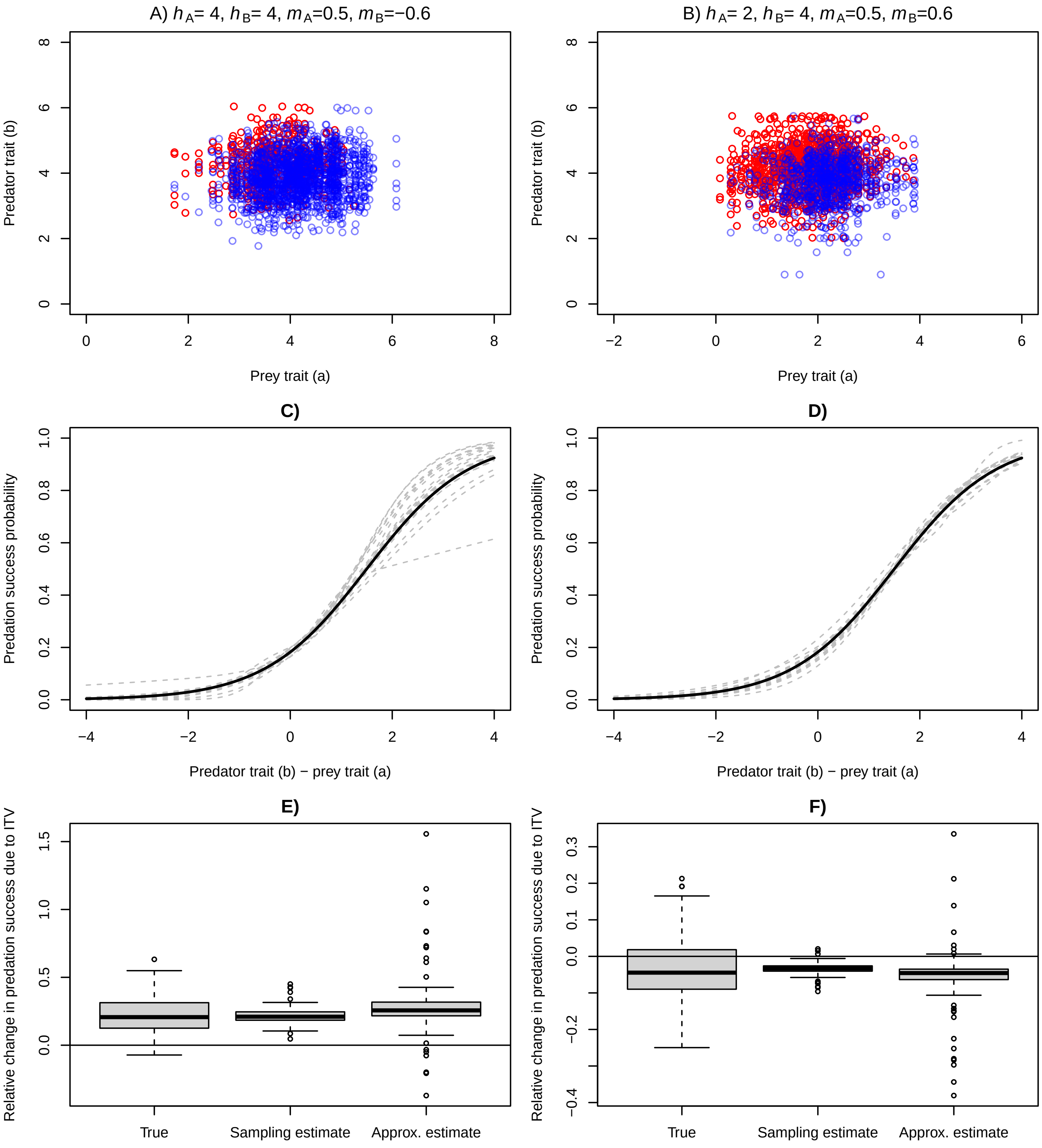
A) hA= 4, hB= 4, mA=0.5, mB=−0.6
B) hA= 2, hB= 4, mA=0.5, mB=0.6
Predator trait (b)
Prey trait (a)
C)
D)
Predation success probability
Predator trait (b) − prey trait (a)
E)
F)
Relative change in predation success due to ITV
True
Sampling estimate
Approx. estimate

**Figure S9.2** – Estimating the impact of ITV on average predation success from field data without environmental variance $\sigma_e = 0$ compared to $\sigma_e = 1.5$ in Fig. 5, and therefore without interspecific trait correlation. The two columns show two scenarios that differ in the reaction norms via which individuals respond to their micro-environment. A, B) Trait values of pairs of interacting individuals in the first replicate, with successful kills shown in red and encounters where the prey escaped shown in blue. Across all replicates in A, there were on average 2179.4 encounters, with 482.6 kills and 1696.8 escapes. In B, there were on average 2186.5 encounters, with 1313.8 kills and 872.7 escapes. C, D) Estimated interaction function for the first 20 replicates shown in dashed gray lines. Note that we have evaluated the two-dimensional interaction function at the respective mean prey trait $h_A$. The true interaction function is shown in black. E, F) Boxplots of true and estimated effects of ITV on total predation success.

### S10 Application example 3: Lotka-Volterra competition model

In the main text, we have explored the effects of ITV and trait correlations in a predator-prey model. As a second example, we here consider a Lotka-Volterra competition model with trait-based competition coefficients. For illustrative purposes we focus on symmetric competition, although our approach works equally well for asymmetric competition (see section S10.5):

$$\begin{aligned}\frac{\mathrm{d}N_A}{\mathrm{d}t} &= r_A N_A \left(1 - \frac{\alpha_{AA}N_A + \alpha_{AB}N_B}{K_A}\right)\\ \frac{\mathrm{d}N_B}{\mathrm{d}t} &= r_B N_B \left(1 - \frac{\alpha_{BB}N_B + \alpha_{AB}N_A}{K_B}\right),\end{aligned} \tag{S10.1}$$

with $N_A$ and $N_B$ the abundances of species $A$ and $B$, $r_i$ their intrinsic growth rates, and $K_i$ their carrying capacities. Furthermore, $\alpha_{AA}$ and $\alpha_{BB}$ are the intraspecific interaction parameters, and $\alpha_{AB}$ is the interspecific interaction parameter.

#### S10.1 Equilibria and coexistence

By setting both derivatives in Eq. S10.1 to zero, the 4 equilibria of the system can be found:

$$(N_A^*, N_B^*) = \begin{cases}(0,0)\\ (\frac{K_A}{\alpha_{AA}}, 0)\\ (0, \frac{K_B}{\alpha_{BB}})\\ (\frac{\alpha_{BB}K_A - \alpha_{AB}K_B}{\alpha_{AA}\alpha_{BB} - \alpha_{AB}^2}, \frac{\alpha_{AA}K_B - \alpha_{AB}K_A}{\alpha_{AA}\alpha_{BB} - \alpha_{AB}^2})\end{cases} \tag{S10.2}$$

Coexistence is possible when both species can invade a system in which only the other species is present, that is, if species $B$ has a positive per-capita growth rate at the equilibrium with only species $A$ $(\frac{K_A}{\alpha_{AA}}, 0)$, and species $A$ has a positive per-capita growth rate in the equilibrium $(0, \frac{K_B}{\alpha_{BB}})$. Therefore, coexistence is possible when the conditions:

$$r_A \left(1 - \frac{\alpha_{AB}K_B}{\alpha_{BB}K_A}\right) > 0 \tag{S10.3}$$

$$r_B \left(1 - \frac{\alpha_{AB}K_A}{\alpha_{AA}K_B}\right) > 0, \tag{S10.4}$$

are both met. Since all parameters are positive, the condition for coexistence simplifies to $\frac{\alpha_{AB}}{\alpha_{BB}} < \frac{K_A}{K_B}$ and $\frac{\alpha_{AB}}{\alpha_{AA}} < \frac{K_B}{K_A}$. When one condition is met, but the other is not, only one of the species will be maintained. In particular, only species $B$ is maintained when $\frac{\alpha_{AB}}{\alpha_{BB}} > \frac{K_A}{K_B}$ and $\frac{\alpha_{AB}}{\alpha_{AA}} < \frac{K_B}{K_A}$. Finally, when neither condition is met, only one of the species will persist depending upon intrinsic growth rates and starting densities. In the special case when carrying capacities are equal, $K_A = K_B$, coexistence will occur when $\alpha_{AB} < \min(\alpha_{AA}, \alpha_{BB})$ i.e. interspecific competition needs to be less than intraspecific competition for coexistence.

#### S10.2 Quantifying the effect of trait variation and correlations

One or more of the competition coefficients can be affected by trait variation, each by averaging their associated interaction function ($\overline{\gamma_{AB}}$, $\overline{\gamma_{AA}}$, and $\overline{\gamma_{BB}}$). Although the shape of the three interaction functions can be chosen independently, the effects of ITV on the average competition coefficients are related through the trait distribution, since the variation in either species affects both average inter- and intraspecific competition.

In the most general model, both intra- and interspecific competition are affected by ITV. Now, each competition coefficient will be calculated using the estimation methods described in section S10.2.1 below, based on their specific interaction functions and trait distributions, and we set $\alpha_{AB} = E[\gamma_{AB}]$, $\alpha_{AA} = E[\gamma_{AA}]$, and $\alpha_{BB} = E[\gamma_{BB}]$. For the intraspecific competition coefficient of species $A$, we denote the joint distribution as $f_{AA}(a, a')$ which is defined as the meeting probability of an individual in species $A$, having phenotype $a$, with another individual of phenotype $a'$ within the same species, with $\rho_{aa'}$ the correlation in trait values across meetings of individuals within the same species. For example, a positive correlation implies that individuals tend to meet with other individuals that have a similar trait value. Although our approach is not restricted to this, for the results in this section we assume a bivariate normal distribution:

$$f_{AA}(a, a') = \frac{1}{2\pi\sigma_a^2\sqrt{1-\rho_{aa'}}} e^{-\frac{1}{2(1-\rho_{aa'})^2}\left[(\frac{a-\mu_a}{\sigma_a})^2 - 2\rho_{aa'}(\frac{a-\mu_a}{\sigma_a})(\frac{a'-\mu_a}{\sigma_a}) + (\frac{a'-\mu_a}{\sigma_a})^2\right]} \tag{S10.5}$$

The joint distribution of meeting probabilities for species $B$ ($f_{BB}(b, b')$) and the interspecific meeting probabilities $f_{AB}(a, b)$ can be defined analogously.

We describe each competition type with a Gaussian interaction function. For interspecific competition this interaction function is defined with parameters $c_{\max,AB}$ and $\omega_{AB}$ as follows:

$$\gamma_{AB}(a, b) = c_{\max,AB} e^{\frac{-(a-b)^2}{\omega_{AB}^2}} \tag{S10.6}$$

Intraspecific interactions in species $A$ are described by the interaction function $\gamma_{AA}(a, a')$, which depends on the trait values $a$ and $a'$ of the two individuals of species $A$ that meet and on the parameters $c_{\max,AA}$ and $\omega_{AA}$ as:

$$\gamma_{AA}(a, a') = c_{\max,AA} e^{\frac{-(a-a')^2}{\omega_{AA}^2}} \tag{S10.7}$$

Similarly, we can define an interaction function for species $B$ denoted by $\gamma_{BB}(b, b')$ as:

$$\gamma_{BB}(b, b') = c_{\max,BB} e^{\frac{-(b-b')^2}{\omega_{BB}^2}} \tag{S10.8}$$

Note that when $\omega_{AA}$ and $\omega_{BB}$ tend to infinity $\alpha_{AA}$ and $\alpha_{BB}$ become equal to $c_{\max,AA}$ and $c_{\max,BB}$ respectively and are no longer trait-dependent. When in addition $c_{\max,AA} = c_{\max,BB} = 1$, the model reduces to a model where intraspecific competition is fixed and only interspecific competition is trait-dependent.

##### S10.2.1 Estimation of average interaction parameters

The naive and exact estimates are obtained for each of the competition kernels separately, in the same way as in the model in the main text. Note, however, that due to the trait means for intraspecific competition being equal (i.e. $\mu_a = \mu_{a'}$ and $\mu_b = \mu_{b'}$), these naive estimates are not trait-dependent:

$$\gamma_{AA}(\mu_a, \mu_a) = c_{\max,AA} \tag{S10.9}$$

$$\gamma_{BB}(\mu_b, \mu_b) = c_{\max,BB}. \tag{S10.10}$$

Using Eq. 10, the second-order Taylor approximations for inter- and intra-specific interaction estimates are:

$$\overline{\gamma_{AB}(a, b)} \approx \gamma_{AB}(\mu_a, \mu_b) + \gamma_{AB}(\mu_a, \mu_b) \cdot \left(\frac{2(\mu_a - \mu_b)^2 - \omega_{AB}^2}{\omega_{AB}^4}\right) \cdot \left(\sigma_a^2 + \sigma_b^2 - 2\sigma_{a,b}\right), \tag{S10.11}$$

$$\overline{\gamma_{AA}(a,a')} \approx c_{\max,AA}\left(1-\frac{2\sigma_a^2}{\omega_{AA}^2}+\frac{2\sigma_{a,a'}}{\omega_{AA}^2}\right), \tag{S10.12}$$

$$\overline{\gamma_{BB}(b,b')} \approx c_{\max,BB}\left(1-\frac{2\sigma_b^2}{\omega_{BB}^2}+\frac{2\sigma_{b,b'}}{\omega_{BB}^2}\right). \tag{S10.13}$$

Similar to the naive estimates, in these Taylor approximations, $\overline{\gamma_{AB}(a,b)}$ is dependent on the difference in trait means, whereas $\overline{\gamma_{AA}(a,a')}$ and $\overline{\gamma_{BB}(b,b')}$ do not depend on trait means.

ITV always decreases the intraspecific interaction parameters $\overline{\gamma_{AA}(a,a')}$ and $\overline{\gamma_{BB}(b,b')}$, whereas within species covariance (positive) always increases them. Note that for some combinations of variance, covariance, mean difference, and $\omega$ the second order Taylor approximation of inter- and intraspecific interaction parameters can become negative. Such regions are biologically infeasible and are not considered for coexistence or population dynamics.

#### S10.3 Results for the case where ITV affects only interspecific competition

We first focus on the case where only interspecific competition is affected by ITV by setting $\alpha_{AA} = \alpha_{BB} = 1$ and $\alpha_{AB} = E[\gamma_{AB}(a,b)] =: \alpha$. The coexistence criterium now reduces to (see also Begon *et al.*, 2006)

$$\alpha < \min\left(\frac{K_A}{K_B}, \frac{K_B}{K_A}\right). \tag{S10.14}$$

The maximal value of this threshold is 1, and occurs when the species' carrying capacities are equal. A necessary, but not sufficient condition for coexistence is thus $\alpha < 1$, which occurs when intraspecific competition is stronger than interspecifc competition, a classical result.

When interspecific competition ($\alpha$) is described by a Gaussian interaction function, trait variation can either increase or decrease average competition, depending on the trait means, as shown in figure S10.1A. Close to the blue diamond, the absolute difference in species' mean trait values is smaller than $\omega/\sqrt{2}$. Here the curvatures are negative and the naive estimate (grey line) is always higher than the estimates that take ITV into account (yellow dots for the approximation and purple and turquoise crosses for the exact estimates). Whether the effect is strong enough to promote coexistence further depends on the value of the coexistence threshold (orange and red lines and diamonds in figure S10.1). When the threshold is slightly below the naive estimate, effects of trait variation can push the average competition below the threshold and foster coexistence (orange threshold, figure S10.1B,D). On the other hand, if the difference in species' trait means is larger than $\omega/\sqrt{2}$ (green diamond), ITV increases the average competition between the species. When the coexistence threshold (red line in figure S10.1A) is just above the naive value, it can hence hinder coexistence (e.g. figure S10.1B,E). Not only qualitative changes (figure S10.1B), but also quantitative changes in equilibrium densities emerge through ITV (e.g. figure S10.1C).

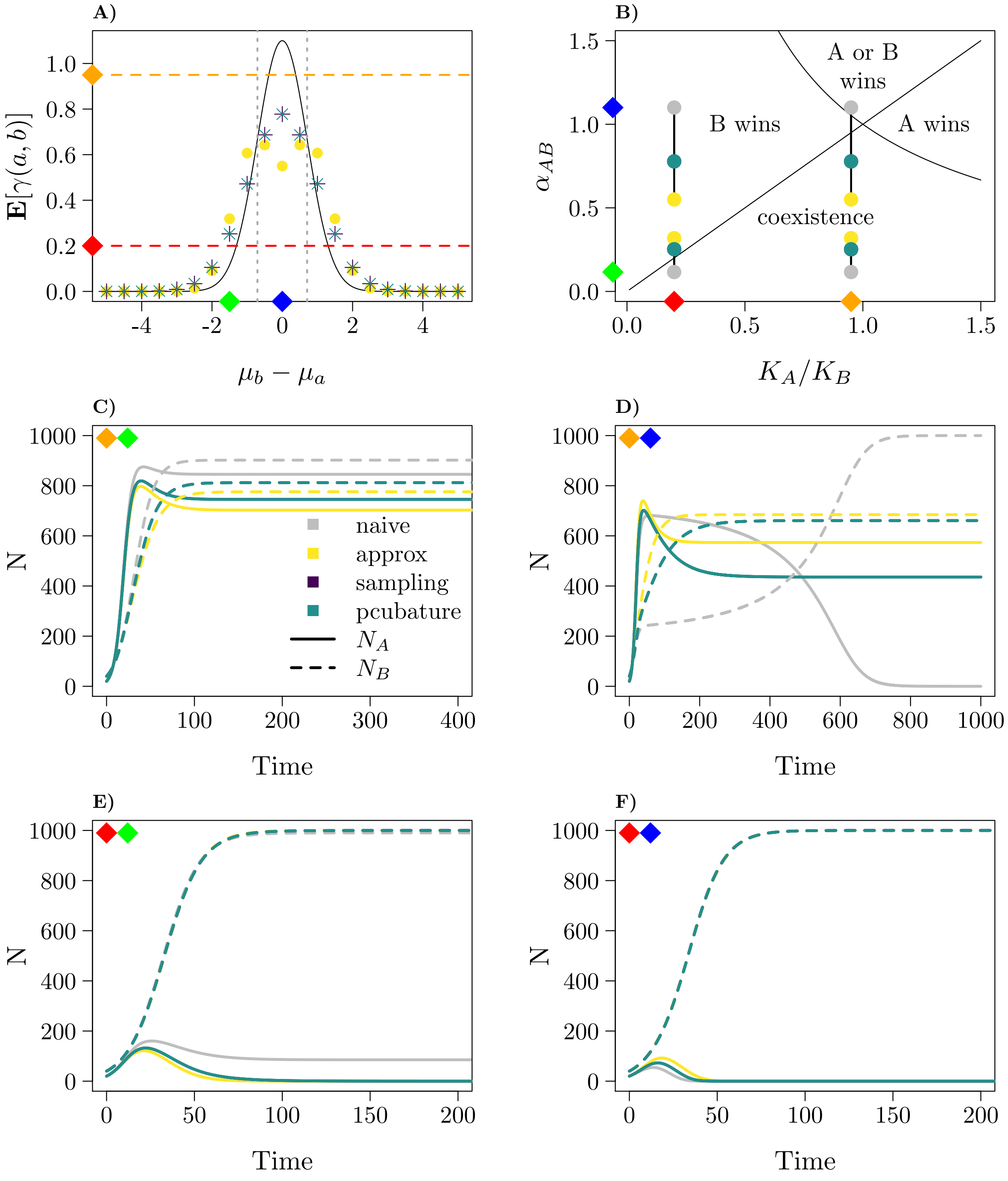
A)
$\mathbf{E}[\gamma(a,b)]$
$\mu_b - \mu_a$
B)
A or B wins
B wins
A wins
coexistence
$\alpha_{AB}$
$K_A/K_B$
C)
naive
approx
sampling
pcubature
$N_A$
$N_B$
N
Time
D)
N
Time
E)
N
Time
F)
N
Time

**Figure S10.1** – Effect of ITV on the dynamics of the Lotka-Volterra competition model when only interspecific competition is trait-dependent. A) Effect of ITV on the average value of a Gaussian interaction function ($c_{\max} = 1.1$, $\omega = 1$). Crosses and points indicate the average interaction function when $\sigma_A = \sigma_B = 0.5$ and $\sigma_{a,b} = 0$ (turquoise, purple, and yellow for the pcubature (numerical integration), sampling, and approximation method respectively). The orange and red lines indicate two possible coexistence thresholds (Eq. S10.14) for $K_A = 950$ and $K_A = 200$, respectively. The green and blue diamonds indicate two trait difference values that are explored in detail in the following panels. The dotted vertical lines indicate $\pm\omega/\sqrt{2}$, the points where the second derivative vanishes. B) Parameter space plot. The different regions are demarcated by the lines $\alpha = K_A/K_B$ and $\alpha = 1/(K_A/K_B)$ C–F) Example time series. The diamonds in the upper left corner indicate the underlying trait difference and threshold value ($K_A/K_B$). Plots for sampling are generally overlaid by pcubature for all the panels. Other parameters: $r_A = 0.2, r_B = 0.1, K_B = 1000$.

#### S10.4 Results for the case where ITV affects both interspecific and intraspecific competition

First, to understand the effect of ITV on intraspecific competition strength, consider the bar plots on the diagonal in figure 3J, which correspond to the average interaction strength between two populations with the same average trait value, as is the case for intraspecific competition. Since the changes are negative compared to the case without ITV, we can see that intraspecific competition strength decreases with increasing trait variance. The reduction in intraspecific competition is minimal when the intraspecific trait correlation is positive and amplified when the intraspecific trait correlation is negative. With normally distributed traits and a Gaussian interaction function, this can also be derived analytically. Let $a$ and $a'$ be drawn from a bivariate normal distribution with mean 0 and variances $\sigma_a^2 = \sigma_{a'}^2 = \sigma^2$ and correlation coefficient $\rho_{aa'}$. Then $\Delta = a - a'$ is also normally distributed with mean 0 and variance $\sigma_\Delta^2 = 2\sigma^2 \cdot (1 - \rho_{aa'})$ (Pishro-Nik, 2014). Integrating over the interaction function then gives

$$\overline{\gamma_{AA}} = \int_{-\infty}^{\infty} \frac{c_{max}}{\sigma_\Delta\sqrt{2\pi}} \cdot e^{-\left(\frac{1}{\omega^2}+\frac{1}{2\sigma_\Delta^2}\right)\Delta^2} d\Delta. \tag{S10.15}$$

With $\sigma' = \sqrt{1/(2/\omega^2 + 1/\sigma_\Delta^2)}$, we obtain

$$\overline{\gamma_{AA}} = \frac{c_{max}}{\sigma_\Delta}\sqrt{\frac{1}{\frac{2}{\omega^2}+\frac{1}{\sigma_\Delta^2}}} \int_{-\infty}^{\infty} \frac{1}{\sigma'\sqrt{2\pi}} e^{-\frac{\Delta^2}{2\sigma'^2}} d\Delta = c_{max}\sqrt{\frac{1}{\frac{4\sigma^2(1-\rho_{aa'})}{\omega^2}+1}}, \tag{S10.16}$$

which clearly decreases with trait variance $\sigma^2$, an effect that is weakened when $\rho_{aa'}$ is positive and amplified when it is negative.

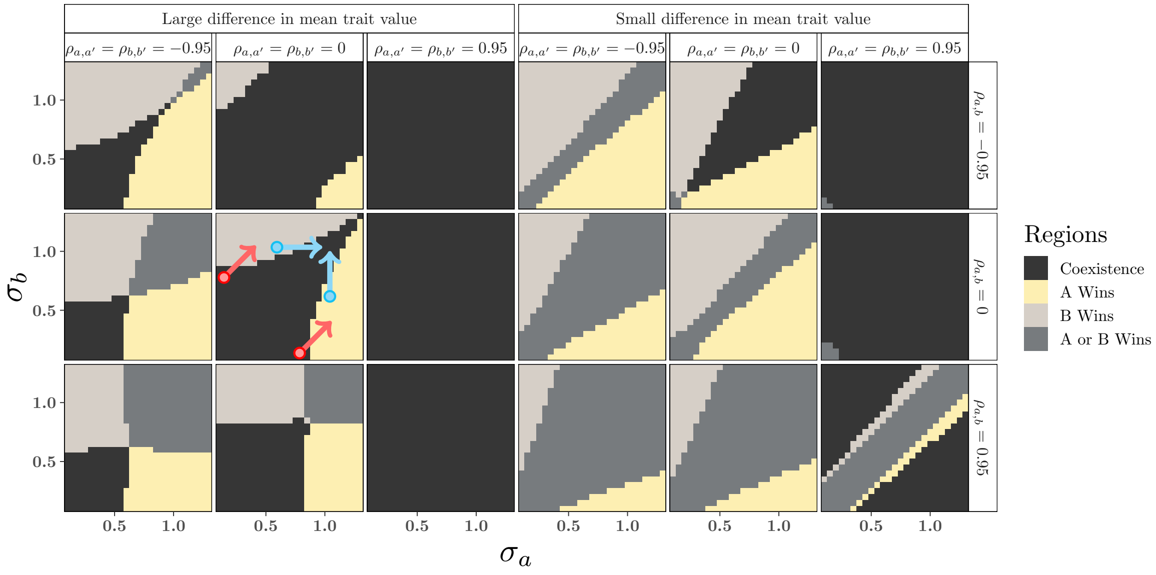

**Figure S10.2** – Effect of ITV ($\sigma_a$ and $\sigma_b$) on coexistence regions for three different inter- and intraspecific trait correlation strengths in the competition models with trait-dependent inter- and intraspecific competition. All $\alpha$s (i.e. $\alpha_{AB}$, $\alpha_{AA}$, and $\alpha_{BB}$) are ITV-dependent. The maximum intraspecific competition coefficients ($c_{\max,AA}$ for species $A$ and $c_{\max,BB}$ for species $B$) are set to 1. Furthermore, all the $\omega$s (i.e. $\omega_{AB}$, $\omega_{AA}$, and $\omega_{BB}$) are equal to 1. Other parameters: maximum interspecific interaction coefficient $c_{\max,AB} = 1.1$, $K_A = K_B = 500$, $\sigma_{a,b} = 0$, $|\mu_a - \mu_b| = 0.9$ for the panels on the left, and $|\mu_a - \mu_b| = 0.2$ for those on the right. Arrows illustrate example scenarios where increases in trait variation either hinder (red arrows) or promote (cyan arrows) coexistence.

The effects of trait variation and intraspecific as well as interspecific trait correlations between interacting individuals on the qualitative outcome of the interaction are explored in figure S10.2. Here we use the sampling method to determine the average interaction parameters, because even small mistakes in approximating the three average interaction parameters can drastically affect their relative strength and thereby lead to faulty prediction of coexistence. Because coexistence is more likely under high intraspecific competition, we generally see larger coexistence regions at high intraspecific trait correlation (figure S10.2).

When the species' trait means are close together (figure S10.2, right half), the effects of ITV on inter- and intraspecific competition are similar: thus interspecific competition also decreases with increasing ITV, with stronger effects when the interspecific correlation coefficient is low (negative, see also figure 3K–L, and Barabás & D'Andrea 2016). Therefore, when the intraspecific trait correlation is larger than the interspecific trait correlation (figure S10.2, top right), an increase in ITV reduces the interspecific competition more than the intraspecific competition, thereby promoting coexistence. In contrast, when all trait correlation coefficients are high, an increase in ITV can only promote coexistence when the increase leads to a larger difference in ITV between the two species (figure S10.2, bottom right). In this case, due to the positive trait correlations, individuals are always matched to individuals with similar trait values regardless of the amount of ITV, and thus competition remains relatively strong. However, due to the difference in trait variation between the species, the most extreme individuals, when matched are still relatively dissimilar, leading to slightly weaker interspecific competition, thereby promoting coexistence. Finally, when trait means are close, ITV in either species can also promote coexistence when interspecific competition has a narrower interaction width compared to intraspecific competition (figure S10.3).

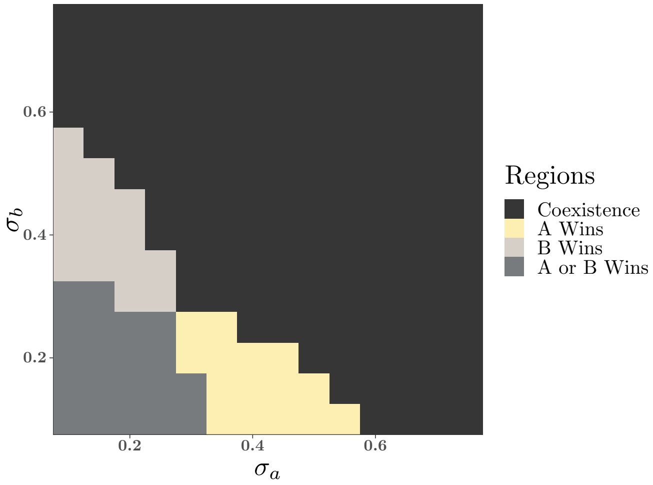

**Figure S10.3** – An example of ITV-dependent inter- and intraspecific competition where higher variance in either of the species promotes coexistence. Here, $\omega_{AB} = 0.4$, $\omega_{AA} = \omega_{BB} = 1$, $c_{\max,AB} = 1.5$, $c_{\max,AA} = c_{\max,BB} = 1$, $|\mu_a - \mu_b| = 0.05$, $\frac{K_A}{K_B} = 1$, and the sampling method is used for estimation of mean interaction.

When the trait means are further apart (figure S10.2, left half), an increase in ITV tends to increase interspecific competition and thereby hinders coexistence (red arrows). The effect is particularly strong when the intraspecific trait correlation is negative, further reducing intraspecific competition. However, even when the means are further apart, ITV can promote coexistence, for example in the absence of trait correlations, if ITV increases in the less variable species (see figure S10.2, blue arrows).

For the case of zero correlations and $K_A = K_B$, figure S10.4 explores more closely how the impact of ITV depends on the $\omega$'s, $c_{\max}$'s, and $|\mu_a - \mu_b|$ and compares the predictions of the Taylor approximation to those of the sampling approach. The regions in figure S10.4 are calculated from the coexistence condition described in section S10.1. Note that the subplots in each panel of figure S10.4 are symmetric around $\sigma_a = \sigma_b$ barring the mirroring of $A$ wins and $B$ wins regions, where $A$ wins when species $A$ has more variation than species $B$ and vice versa. Moreover, in some subplots coexistence is favoured when both species have similar amounts of trait variation. As a consequence, when the amount of trait variation is much larger in one of the species, coexistence may be favoured by an increase in trait variation in the other species (e.g. second row from the top in the bottom right panel of figure S10.4).

When variance is small, the coexistence predictions obtained from the second order Taylor approximation agree with those based on the sampling estimates. However, as the variances increase, the predictions from the approximation diverge from the exact predictions from the sampling method. This is a consequence of large variation in curvatures over the joint trait distribution. When the difference between trait means is small, curvatures also vary strongly over the distribution, thereby inducing similar inaccuracies in the average competition parameters. However, in this case, due to the means being close, the bias in the three competition parameters is similar, and when comparing them to determine coexistence, the bias partly cancels out. This yields a slightly more accurate approximation of the coexistence region when means are close (figure S10.4, bottom right). For large amounts of ITV, in particular when $\sigma_{ii} > \omega/\sqrt{2}$, the approximation of the intraspecific competition coefficient even becomes negative, as indicated by the red striped regions. For larger $\omega$, these infeasible regions are smaller, due to the scaling effect that $\omega$ has on the trait values (figure S10.4).

When $\frac{\omega}{\sqrt{2}} > |\mu_a - \mu_b|$ (see Eq. S10.11), variance reduces the interspecific competition, and for large enough variance, $\alpha_{AB}$ becomes negative. This causes the infeasible regions in the respective approximation panels with fixed intraspecific competition in figure S10.4. On the other hand, $\alpha_{AB}$ increases with $\sigma$ when $\frac{\omega}{\sqrt{2}} < |\mu_a - \mu_b|$ and hence does not become negative which leads to no infeasible regions in the respective approximation panel of fixed intraspecific competition.

Importantly, when we compare the results for fixed intraspecific competition (column 1, figure S10.4) with variable intraspecific competition (column 2, figure S10.4) we note that the coexistence regions shrink when both intra- and interspecific competition are affected by ITV. Similarly, within any subpanel for the variable intraspecific competition case, coexistence becomes less likely with increasing variances. This is expected behaviour, because in the used set up, interspecific competition is reduced less (and it is potentially even increased) than intraspecific competition. This finding is however dependent on intra- and interspecific competition acting in a similar manner. When competition width for intraspecific competition is much larger than for interspecific competition, the opposite effect can occur (see for an example figure S10.3).

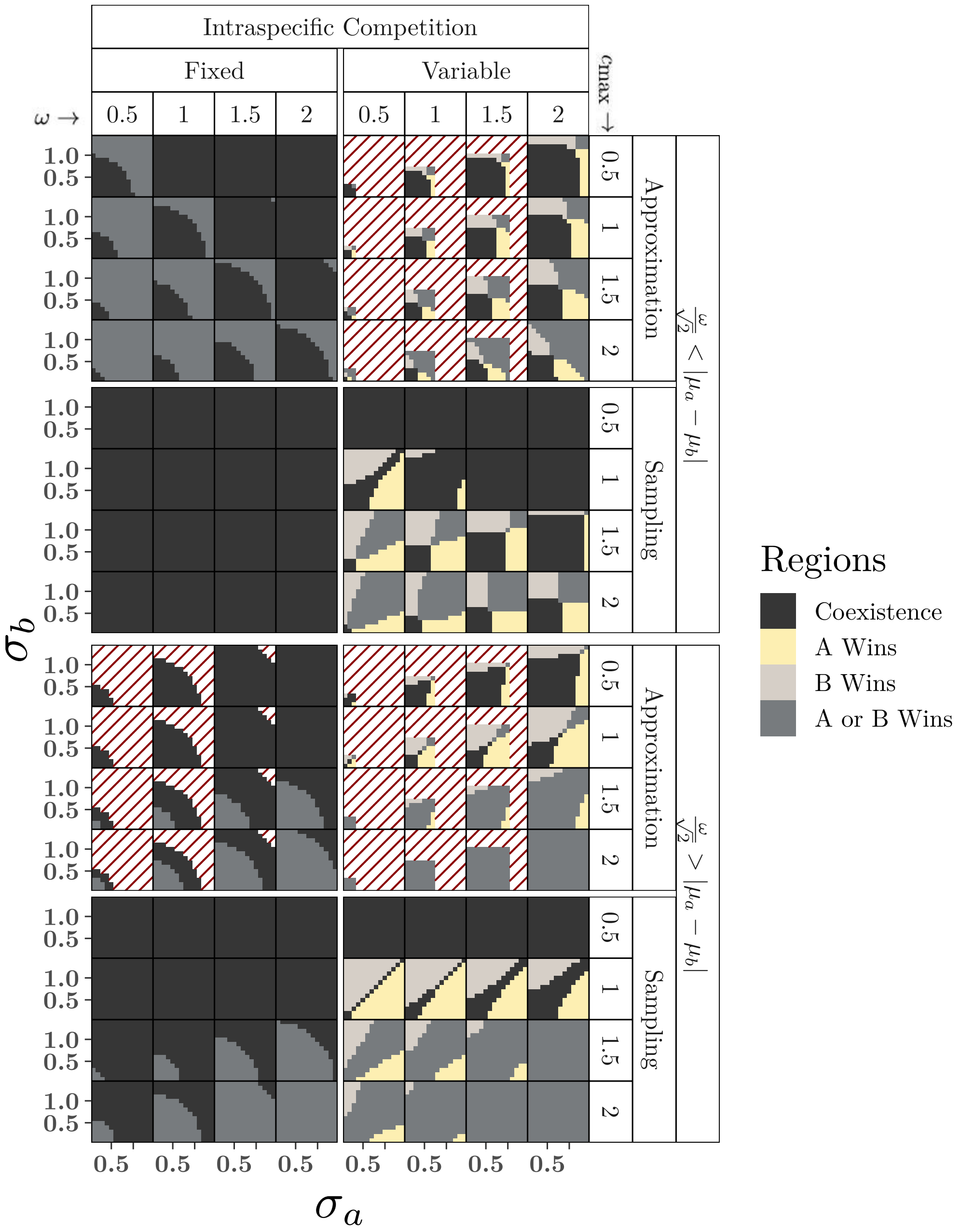

**Figure S10.4** – The effect of ITV ($\sigma_a$ and $\sigma_b$), width ($\omega$), and $c_{\max}$ on regions of coexistence. The red striped area is the infeasible region (i.e. negative values of the average interaction parameter estimated by the approximation). The left column is for fixed intraspecific competition (i.e. $\alpha_{AA} = \alpha_{BB} = 1$), and the right column is for variable intraspecific competition. To ensure comparability between the two columns only the $c_{\max,AB}$ (here $c_{\max}$) is changed, and $c_{\max,AA}$ and $c_{\max,BB}$ are fixed to 1. All the $\omega$ are equal (i.e. $\omega_{AB} = \omega_{AA} = \omega_{BB} = \omega$). $|\mu_a - \mu_b| = 1.5 \cdot \frac{\omega}{\sqrt{2}}$ under the condition $\frac{\omega}{\sqrt{2}} < |\mu_a - \mu_b|$ and $|\mu_a - \mu_b| = 0.5 \cdot \frac{\omega}{\sqrt{2}}$ under the condition $\frac{\omega}{\sqrt{2}} > |\mu_a - \mu_b|$. Carrying capacities $K_A = K_B = 500$, and correlation is kept zero.

#### S10.5 Discussion

Compared to the predator-prey model analysed in the main text, the effect of trait variation and correlations on coexistence is even stronger in the competition model. When ignoring trait correlations, ITV tends to hinder coexistence because it can only decrease intraspecific competition, in agreement with similar work by Hart *et al.* 2016. A change in the trait correlations can affect coexistence itself, but also whether changes in ITV promote or hamper coexistence (e.g. compare the panel in the bottom right, to the panel on the top row, fifth column in figure S10.2). Trait correlations could be the consequence of spatial structure in the populations. For example, a negative trait correlation corresponds to the case where individuals tend to co-occur more with heterospecific individuals that are relatively dissimilar, and thus reduced similarity-based competition (see figure 3L). This could happen for example because the two competing species show opposite responses to a micro-environmental factor, as observed for some traits in sub-Arctic plants (Kemppinen & Niittynen, 2022, see also section 2.1.1). Spatial structure can profoundly shape the effects of ITV, as also shown by Uriarte & Menge (2018). In their two-patch extension of the model by Hart *et al.* (2016), intraspecific variation can promote coexistence via nonlinear averaging if the overall inferior competitor is favored in the more productive patch, the opposite of what is found in the non-spatial model. However, in a model with evolution of trait means and variances, Wickman *et al.* (2023) found that the number of stably coexisting species was generally lower with evolving ITV than without ITV, both with or without spatial structure. Although nonlinear averaging over the joint trait distribution can capture some potential effects of spatial structure, it does not affect the degrees of freedom of the population model and it can therefore not produce the full dynamics that have been found in models that account for space more explicitly (see e.g. Bascompte & Solé, 1995). In comparison, we expect our model to describe a limiting case where the spatial distribution equilibrates over short time scales relative to the time scale of population dynamics. Our approach can then be used to obtain a first intuition when not enough information is available for a spatially explicit model.

These examples show that a full understanding of the population dynamics require simultaneously taking the variation in both species and their trait correlations into account. For competition models, Stump *et al.* (2022) recently synthesized how ITV affects coexistence via its influences on fitness differences between species and stabilizing mechanisms. They find that in large part the results depend on whether the variable traits are "niche traits" (for which performance can vary with the conditions, e.g. types of food available) or "hierarchical traits" (for which the rank order of performance of different values is independent of the conditions). Here we focused on a Gaussian matching trait function for the competition scenario, such that the relative performance of trait values in one species would depend on the trait distribution in the other species, making the trait a niche trait *sensu* Stump *et al.* (2022). For such niche traits, they find that an increase in ITV can enhance the stabilizing effect and thus coexistence, if ITV is increased more in the more variable species, thereby creating a generalist-specialist trade-off. We also observe this in Fig. S10.2. However, as described above, we observe a number of other ways in which ITV can promote coexistence, which are mediated by individual-by-individual trait correlations and thus, as far as we understand, are not captured by the framework of Stump *et al.* (2022).

#### References in the SI

Barabás, G. & D'Andrea, R. (2016) The effect of intraspecific variation and heritability on community pattern and robustness. *Ecology Letters* **19**, 977–986.

Bascompte, J. & Solé, R.V. (1995) Rethinking complexity: modelling spatiotemporal dynamics in ecology. *Trends in Ecology & Evolution* **10**, 361–366.

Begon, M., Townsend, C.R. & Harper, J.L. (2006) *Ecology - From Individuals to Ecosystems*. Blackwell Publishing Ltd, Oxford, UK, 4th edn.

Gibert, J.P. & Brassil, C.E. (2014) Individual phenotypic variation reduces interaction strengths in a consumer-resource system. *Ecology and Evolution* **4**, 3703–3713.

Hart, S.P., Schreiber, S.J. & Levine, J.M. (2016) How variation between individuals affects species coexistence. *Ecology Letters* **19**, 825–838.

Isserlis, L. (1918) On a formula for the product-moment coefficient of any order of a normal frequency distribution in any number of variables. *Biometrika* **12**, 134–139.

Kemppinen, J. & Niittynen, P. (2022) Microclimate relationships of intraspecific trait variation in sub-arctic plants. *Oikos* **2022**, e09507.

Narasimhan, B., Johnson, S.G., Hahn, T., Bouvier, A. & Kiêu, K. (2019) *cubature: Adaptive Multivariate Integration over Hypercubes*. R package version 2.0.4.

Phillips, B.L., Brown, G.P. & Shine, R. (2003) Assessing the potential impact of cane toads on Australian snakes. *Conservation Biology* **17**, 1738–1747.

Phillips, B.L. & Shine, R. (2004) Adapting to an invasive species: toxic cane toads induce morphological change in Australian snakes. *Proceedings of the National Academy of Sciences of the United States of America* **101**, 17150–17155.

Pishro-Nik, H. (2014) *Introduction to probability, statistics, and random processes*. Kappa Research LLC.

Senthilnathan, A. & Gavrilets, S. (2021) Ecological consequences of intraspecific variation in coevolutionary systems. *The American Naturalist* **197**, 1–17.

Stump, S.M., Song, C., Saavedra, S., Levine, J.M. & Vasseur, D.A. (2022) Synthesizing the effects of individual-level variation on coexistence. *Ecological Monographs* **92**, e01493.

Uriarte, M. & Menge, D. (2018) Variation between individuals fosters regional species coexistence. *Ecology Letters* **21**, 1496–1504.

Wickman, J., Koffel, T. & Klausmeier, C.A. (2023) A theoretical framework for trait-based eco-evolutionary dynamics: Population structure, intraspecific variation, and community assembly. *The American Naturalist* **201**, 501–522.
